# Pruning the Search, Not the Signal: Adaptive-Banding Needleman–Wunsch Sequence Alignment via Protein Language Model Confidence

**DOI:** 10.64898/2026.08.26.747234

**Authors:** Muhammad Shoaib, Waqas Ali

**Author notes:** Corresponding author. *Email addresses:* (Muhammad Shoaib), (Waqas Ali).

## Abstract

Dynamic programming (DP) yields exact quadratic-time (O(*NM*)) pairwise sequence alignments. Static banding heuristics (O(*NW*)) fail catastrophically on low-identity (< 30%), asymmetric insertions/deletions (indels), or extreme length ratios, dropping core-block Sum-of-Pairs (SP) score recovery to 20%–50%. Conversely, recent protein language model (PLM) aligners evaluate all *N* × *M* cells without search grid constraints. To bridge this gap, we introduce Adaptive-Banding Needleman–Wunsch (AB-NW), pruning the search space without sacrificing sequence alignment signal by leveraging PLM contextual representations to construct a confidence-adaptive DP corridor prior to fine-resolution DP while keeping downstream scoring unmodified. AB-NW downsamples residue embeddings, computes a coarse alignment, and sets per-row corridor bounds via normalized confidence metrics. Evaluated via JIT-compiled buffers, this reduces time complexity to 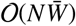 and space to O(*NW*_max_), where 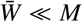. Benchmarked across three PLM backbones (`ESM2-8M, ESM2-35M, ProtBERT`) across nine structural challenge categories, AB-NW recovers > 98.9% of exact unconstrained alignment scores and core-block SP accuracy across static banding failure modes (Twilight Zone, Asymmetric Indels, Extreme Aspect Ratios) while eliminating 55.3%–78.8% of active DP cells. On large protein sequence matrices (*N, M* ≥ 3,700), AB-NW eliminates 87.6%–91.7% of cells, achieving speedups of 9.79×–13.30× (pure DP) and 1.73×–2.94× (end-to-end), reaching up to 18.12× on unbiased controls (*p* < 0.05 to *p* < 10^−15^), making AB-NW practical for large-scale, high-throughput sequence alignment pipelines.

**Graphical Abstract:** 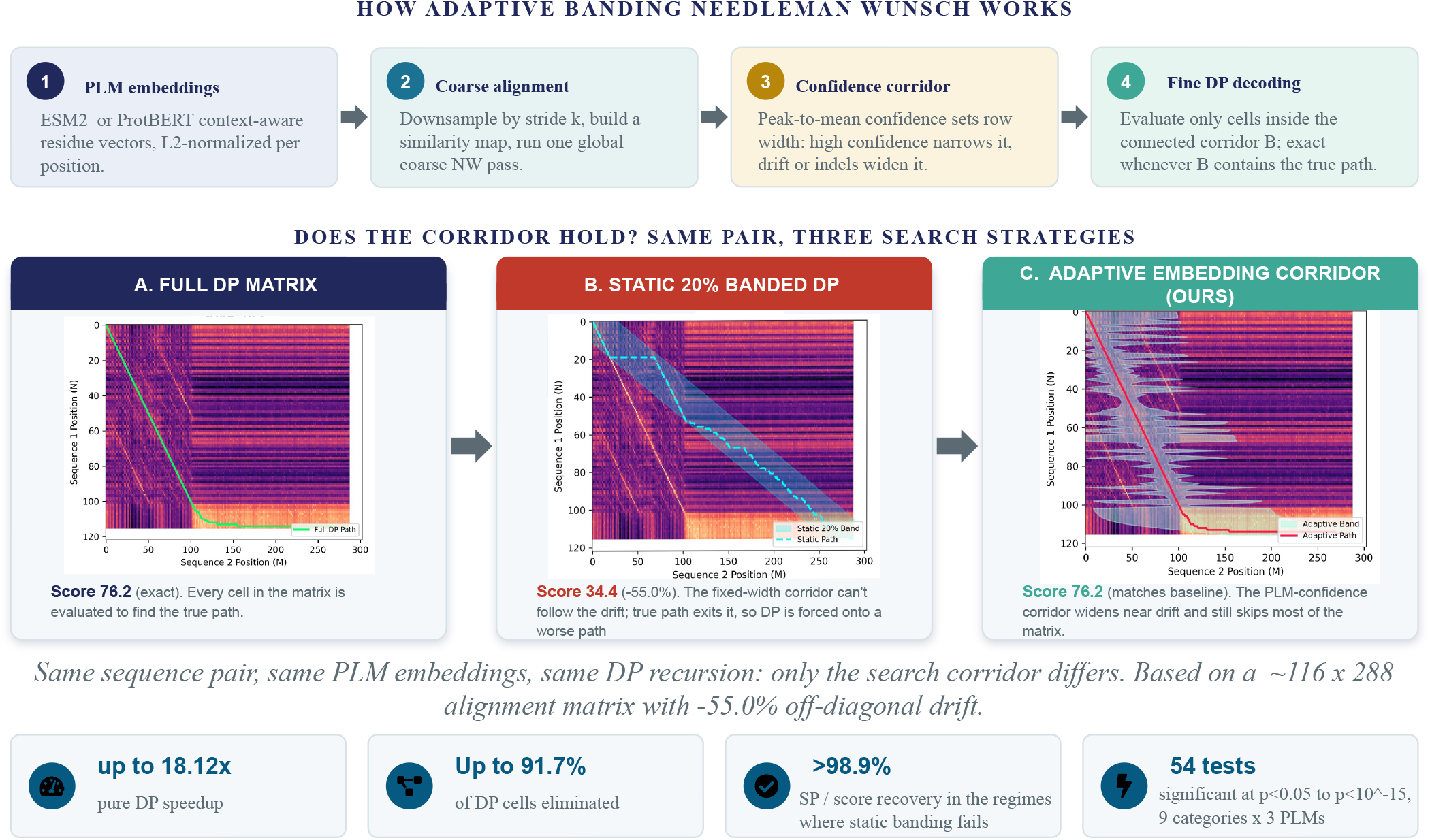

## 1. Introduction

Sequence databases continue to grow rapidly: UniProtKB alone holds over 250 million protein sequences [1], and raw sequence data offers limited utility until systematically compared against known targets. Pairwise sequence alignment underpins computational biology, driving homology inference, functional annotation, phylogenetics, and multiple sequence alignments (MSAs) that supply co-evolutionary signals to structure predictors like AlphaFold [2]. Alignment quality directly dictates downstream model performance on representation-learning benchmarks like TAPE [3] and zero-shot mutation-effect predictions by PLMs [4]. Indeed, large-scale fitness evaluations in ProteinGym [5] demonstrate that alignment variance, rather than model architecture, explains reported benchmark performance differences. Producing accurate alignments efficiently at scale remains a non-trivial challenge.

Scalable bioinformatic workflows rely on heuristic searchers like BLAST [6], MMseqs2 [7], and Foldseek [8]. While indexing exact sequence or structural seeds maximizes query throughput, these utilities cannot compute exact residue-to-residue alignments for detailed pairwise comparisons.

Exact global alignment relies on Needleman–Wunsch dynamic programming (DP) [9], extended by Gotoh to affine gap penalties [10]. Standard DP requires O(*NM*) time and space for an *N* × *M* matrix, stalling large proteomic workflows. Static or score-driven banding mitigates this burden to O(*NW*) by confining evaluation to a corridor around diagonal *j* = *i* · *M/N* [11]. However, within the sequence identity twilight zone of 20%– 35%, or amid severe length asymmetry and insertions, static corridors clip the optimal path, causing severe accuracy drops [12].

Protein language models (PLMs) like ESM [13] and Prot-BERT [14] generate contextual representations that preserve structural homology at low sequence identities. Modern aligners exploit these embeddings to improve fidelity, including EBA [15], PEbA [16], and pLM-BLAST [17]. Yet these techniques inject embeddings into cell scoring rather than search-grid pruning. Because they still compute the complete O(*NM*) matrix or rely on static diagonal windows, quadratic computational complexity remains unaddressed. This gap frames three core questions:

1. RQ1: Can precomputed PLM embedding confidence guide dynamic programming cell selection as an alternative to fixed or score-driven corridors, without modifying the underlying scoring metric?
2. RQ2: Under what structural regimes (e.g., twilight-zone identities, large or asymmetric indels, and extreme length ratios) does a confidence-bounded corridor preserve reference accuracy while skipping most cell evaluations?
3. RQ3: Does this search corridor generalize across varied PLM architectures, embedding dimensions, and dynamic programming recursions?

To address these questions, we present AB-NW, a framework that uses PLM representations to construct a confidence-adaptive search corridor prior to fine-resolution DP without altering downstream scoring functions. AB-NW aligns downsampled residue embeddings globally and evaluates row-wise confidence signals to vary corridor width dynamically. The corridor contracts across high-confidence regions and expands across structural perturbations such as indels. Decoupling corridor construction from intermediate DP scores eliminates runtime recentering overhead, generalizing directly across global, local, and affine-gap recursions while lowering complexity to 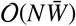.

We evaluate AB-NW against exact unconstrained DP across six static-band baselines and nine benchmark regimes: twilight-zone pairs, ultra-giant matrices, diverse large matrices, large asymmetric indels, extreme length ratios, internal repeats, micro-peptides, standard-length pairs, and an unbiased control set. The test suites derive from a family-level, leakage-free partition of BAliBASE, PREFAB, OXBench, and SABmark [18, 19, 20, 21]. Hyperparameters were tuned across three PLM backbones of disparate depth and topology via Tree-structured Parzen Estimator (TPE) Bayesian optimization [22]. Where static banding collapses to 20%–50% recovery on twilight-zone pairs and large indels, AB-NW retains over 99% of reference accuracy while bypassing most DP cell evaluations across all regimes.

Principal contributions include:

1. Decoupled corridor construction: Unlike EBA, PEbA, and pLM-BLAST, AB-NW applies embedding similarity exclusively to restrict DP search boundaries upfront, retaining complete compatibility with downstream substitution matrices and scoring models.
2. Local adaptive confidence metrics: A peak-to-mean confidence metric derives row-wise corridor bounds locally instead of imposing a uniform width. Banded decoding is proven to recover the exact global optimum whenever the true path falls within the corridor (Proposition 1), bounding runtime to 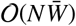 and memory to O(*NW*_max_) (Proposition 2).
3. Recursion-agnostic formulation: Coarse-to-fine corridor construction transfers directly to global (Needleman– Wunsch), local (Smith–Waterman), and affine-gap (Gotoh) recursions without structural modifications or separate tuning.
4. Leakage-free multi-regime evaluation: Testing spans a strict family-level split across four established databases, partitioned into nine regimes designed around static banding failure points.
5. Open, reproducible implementation: JIT-compiled CPU and GPU routines execute corridor generation and banded DP. A Bayesian objective (Eq. (7)) explicitly penalizes performance drops in any individual benchmark regime during hyperparameter search.

The remainder of this paper is organized as follows. Section 2 reviews dynamic programming, static banding heuristics, and PLM aligners. Section 3.1 formalizes adaptive corridor construction, establishes complexity bounds, and extends the framework to local and affine-gap variants. Section 3.8 details the benchmark corpus, family-level partitioning, and hyperparameter optimization. Section 4 reports empirical performance across all nine benchmark regimes and three PLM backbones.

## 2. Literature Review

AB-NW intersects exact dynamic-programming (DP) alignment, search space reduction, and protein language model (PLM) representations. Prior PLM aligners substitute embedding similarity into the *scoring* function while evaluating the full *N* × *M* grid. AB-NW instead preserves the recursion untouched, employing embeddings strictly to restrict evaluated cells.

### 2.1. Exact Dynamic-Programming Alignment

Needleman and Wunsch [9] formulated global alignment over an (*N*+1) × (*M*+1) matrix in O(*NM*) time and space. Smith and Waterman [23] adapted this recursion to local alignment by flooring cell scores at zero; Gotoh [10] added affine gap costs at a constant factor memory increase. AB-NW resolves this shared quadratic bottleneck; Section 3.5 shows all three recursions decode unchanged within an adaptive corridor.

Space optimizations historically targeted linear bounds. Hirschberg [24] achieved O(min(*N, M*)) space via divide and conquer midpoint crossings; Myers and Miller [25] paired this with Gotoh’s affine recursion to obtain O(*N*+*M*) space at roughly double the runtime. AB-NW instead retains active corridor cells in row compressed buffers, securing O(*NW*_max_) space without recomputation overhead. Other accelerations exploit input structure. Myers’ O(*ND*) difference algorithm [26] sweeps diagonally and scales near linearly for closely related sequences, while bit vector parallelism [27] evaluates cell differences rather than scores. Both assume unit cost discrete edits and fail on continuous embedding similarities. Classical DP scoring tables, PAM [28] and BLOSUM [29], assign context free scores to residue pairs and degrade below ∼50% sequence identity. AB-NW remains agnostic to matrix choice; any substitution table decodes within the corridor.

### 2.2. Restricting the Search Space: Static and Adaptive Banding

Static banded DP evaluates a corridor of width *w* = *f* · *M* along diagonal *j* = *i* · *M/N* [30], reducing cost to O(*NW*). The band succeeds only when optimal paths track the diagonal closely. Indels and sequence length asymmetries displace paths outward; on divergent pairs (Categories 1, 4, and 5), fixed 5% bands recover only 20 to 30% of reference core blocks.

Adaptive banding addresses this drift dynamically. Suzuki and Kasahara [11] track antidiagonal score trajectories to recenter fixed width bands on noisy nucleotide reads; ABSW and GACT stitch fixed size overlapping windows, while Nanopolish ABEA applies banded DP to raw nanopore signals. Gao et al. [31] implement SIMD accelerated adaptive limits for partial order alignment graphs. Because these frameworks deduce steering signals from cumulative DP scores, long runs of low scoring gap extensions can truncate bands prematurely. AB-NW diverges on three points: corridor width varies per row, predecoding coarse embedding confidence guides adaptation rather than accumulated scores, and boundaries freeze before fine resolution decoding begins (Section 3.3). This coarse to fine strategy transposes multiresolution dynamic time warping with Sakoe–Chiba bands [32, 33] to residue embeddings.

Unbanded alternatives diverge in design. Wavefront Alignment and its bidirectional extension [34, 35] achieve O(*Ns*) runtime for edit distance *s*, which accelerates close sequences but falters in the low identity, heavily gapped regime targeted here. Parallel hardware reduces cost per cell rather than grid volume: striped SIMD vectorization [36], GPU wavefronts [37], and in memory processing units [38, 39], as well as FPGA-based wavefront accelerators [40] and cross-architecture parallel frameworks for dynamic programming [41]. Operating at a distinct layer, these designs compose directly with corridor restriction.

### 2.3. Protein Language Models and Embedding-Guided Alignment

Masked language model transformers [42] acquire representations capturing secondary structure and contacts without supervision [43], scaling with model capacity [13]. ProtBERT [14] yields a more diffuse embedding space than ESM-2, shaping tuned hyperparameters directly (Table 4).

Embeddings increasingly guide alignment algorithms. EBA [15] and PEbA [16] substitute BLOSUM values with cosine similarities across unbanded NW and Smith–Waterman recursions to elevate twilight zone accuracy. DEDAL [44] learns position specific gap and substitution parameters through differentiable modules, extended to MSA optimization by Petti et al. [45]. PROSTAlign [46] trains a 400 × 400 pair substitution matrix; Ankh-score [47, 48] and OTalign [49] similarly transform all cell scores. Because each evaluates the complete O(*NM*) grid, quadratic space complexity remains. Conversely, pLM-BLAST [17] and the positional embedding search of Johnson et al. [50] restrict grids via seeded static diagonals, inheriting Section 2.2 band failures. Embeddings also bypass residue level DP during homology screening [51] or MSA clustering [52, 53, 54], sacrificing exact pairwise decodings. Nearest conceptually is the coarse to fine double DP of Spicer et al. [55]; AB-NW differs because its coarse pass outputs corridor boundaries and confidence statistics rather than a guiding path, leaving the fine pass an exact, unmodified DP decoder over those bounds. Decoupling corridors from scoring allows any embedding scoring model to execute within AB-NW corridors.

### 2.4. The Sequence-Identity Twilight Zone

Rost [12] showed that sequence identity predicts structural similarity reliably only above ∼40%, failing to discriminate homologous from nonhomologous pairs below ∼35%. This threshold demarcates the Category 1 benchmark (SeqID ≤ 30%, Drift_max_ ≥ 20 residues), where static bands clip true trajectories. Profile HMM methods [56, 57, 58] identify remote homology but require precomputed MSAs; AB-NW evaluates single sequence pairs directly, obtaining comparable contextual sensitivity from pretrained representations.

### 2.5. Positioning of the Present Work

Table 1 organizes the design space across five axes: PLM input, search space restriction, localized band variance, unmodified score recursions, and predecoding restriction independent of accumulated DP values. AB-NW is the sole entry satisfying all five, uniquely combining space restriction with a conditional exactness guarantee (Proposition 1).

**Table 1:** Comparison of DP aligners, banding methods, and PLM-guided alignment across search constraints and asymptotic complexity.

| Method | PLM input | Restricts DP search | Adaptive width | Scoring unmodified | Upfront restriction | Time complexity |
| --- | --- | --- | --- | --- | --- | --- |
| *Exact discrete DP* |  |  |  |  |  |  |
| Needleman–Wunsch [9] | × | × | × | ✓ | — | $O(NM)$ |
| Smith–Waterman [23] | × | × | × | ✓ | — | $O(NM)$ |
| Gotoh (affine) [10] | × | × | × | ✓ | — | $O(NM)$ |
| *Search-space reduction* |  |  |  |  |  |  |
| Static diagonal band [30] | × | ✓ | × | ✓ | ✓ | $O(NW)$ |
| Suzuki–Kasahara [11] | × | ✓ | × | ✓ | × | $O(NW)$ |
| abPOA [31] | × | ✓ | × | ✓ | × | Sub-quad. |
| WFA / BiWFA [34, 35] | × | ✓ | ✓ | × | × | $O(Ns)$ |
| *PLM-guided alignment* |  |  |  |  |  |  |
| vcMSA [52] | ✓ | N/A<sup>c</sup> | N/A | × | N/A | N/A |
| pLM-BLAST [17] | ✓ | ✓ | × | × | ✓ | $O(NW)$ |
| EBA [15] | ✓ | × | × | × | — | $O(NM)$ |
| PEbA [16] | ✓ | × | × | × | — | $O(NM)$ |
| DEDAL [44] | ✓ | × | × | × | — | $O(NM)$ |
| Ankh-score [47] | ✓ | × | × | × | — | $O(NM)$ |
| OTalign [49] | ✓ | × | × | × | — | $O(NM)$ |
| Coarse-to-fine double DP [55] | ✓ | ✓ | × | × | ✓ | $O(NM)$ |
| **AB-NW (this work)** | ✓ | ✓ | ✓ | ✓ | ✓ | $O(N\bar{W})$ |

<sup>a</sup>Band shifts dynamically; width is fixed. <sup>b</sup>Requires edit-distance costs. <sup>c</sup>Non-DP method.
Notation: $N, M$ : sequence lengths; $W, \bar{W}$ : fixed/mean band width ( $\bar{W} \ll M$ ); $s$ : edit distance. Space complexity: $O(NM)$ unconstrained, $O(NW)$ banded, $O(s)$ BiWFA, $O(NW_{\max})$ AB-NW.

## 3. Methodology

### 3.1. Problem Formulation

Needleman–Wunsch (NW) global pairwise alignment of sequences *P* = *p*_1_ *p*_2_ … *p*_*N*_ and *Q* = *q*_1_*q*_2_ … *q*_*M*_ requires evaluating an (*N*+1) × (*M*+1) dynamic-programming (DP) matrix in *O*(*NM*) time and space [9]. Quadratic scaling bottlenecks long sequences. Static banded alignment restricts the DP search space to an ad hoc corridor around the main diagonal, risking optimality if the true path drifts.

We address this limitation by extracting per-residue embeddings from pretrained PLMs to predict row-specific column bounds [*L*_*i*_, *R*_*i*_]. Instead of imposing fixed widths, our method dynamically scales the band via a local confidence signal. The resulting AB-NW algorithm confines exact DP recursion to this data-dependent corridor. This bounds expected time complexity to 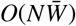 with 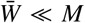 (Proposition 2), preserving exact DP decoding without relying on heuristic path approximations.

#### Definition 1

(Alignment corridor). An alignment corridor for sequences of length *N* and *M* is a family of integer intervals 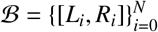, 0 ≤ *L*_*i*_ ≤ *R*_*i*_ ≤ *M*, over which the DP recursion (Eq. (5)) is evaluated. *ℬ* is *connected* if *L*_*i*_ ≤ *R*_*i*−1_ + 1 and *R*_*i*−1_ ≥ *L*_*i*_ − 1 for all *i*, ensuring every in-band cell remains reachable from (0, 0) via in-band diagonal, vertical, or horizontal steps.

The pipeline spans four steps: PLM embedding extraction (Algorithm 1), coarse-to-fine corridor construction (Algorithms 2–3 banded DP decoding (Algorithm 4), and Bayesian hyperparameter optimization (Algorithm 5). Table 2 outlines the notation. Figure 1 details the architecture.

**Table 2:**
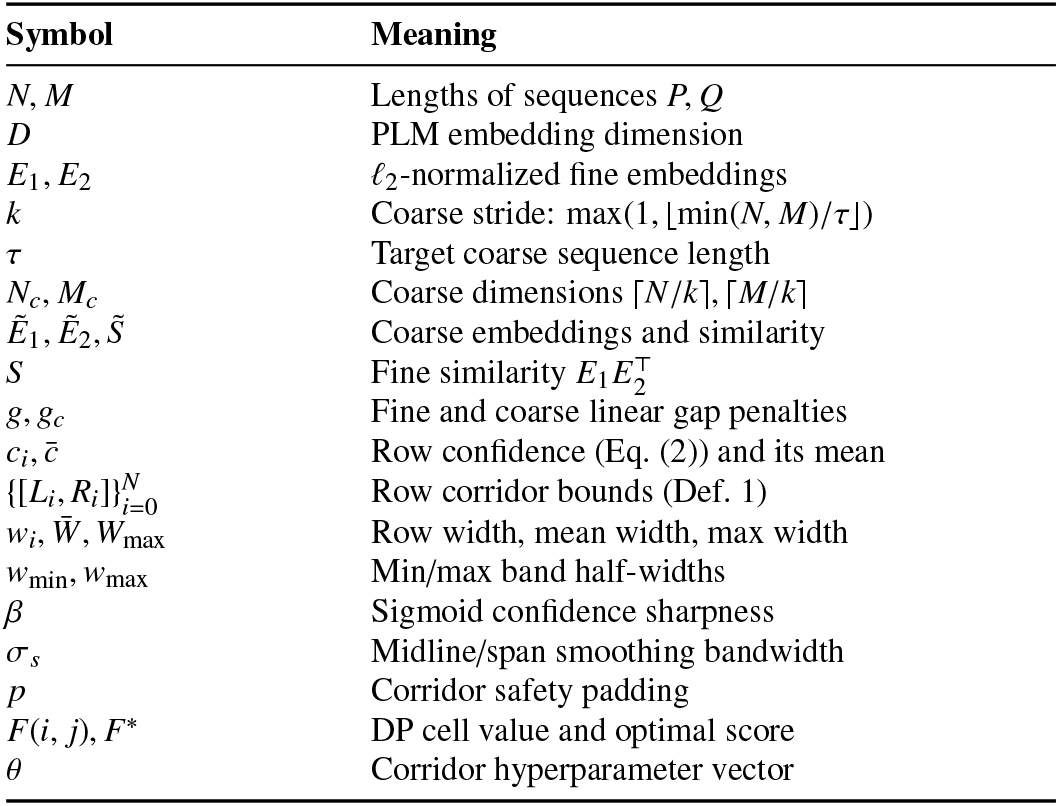
Summary of notation.

**Figure 1:**
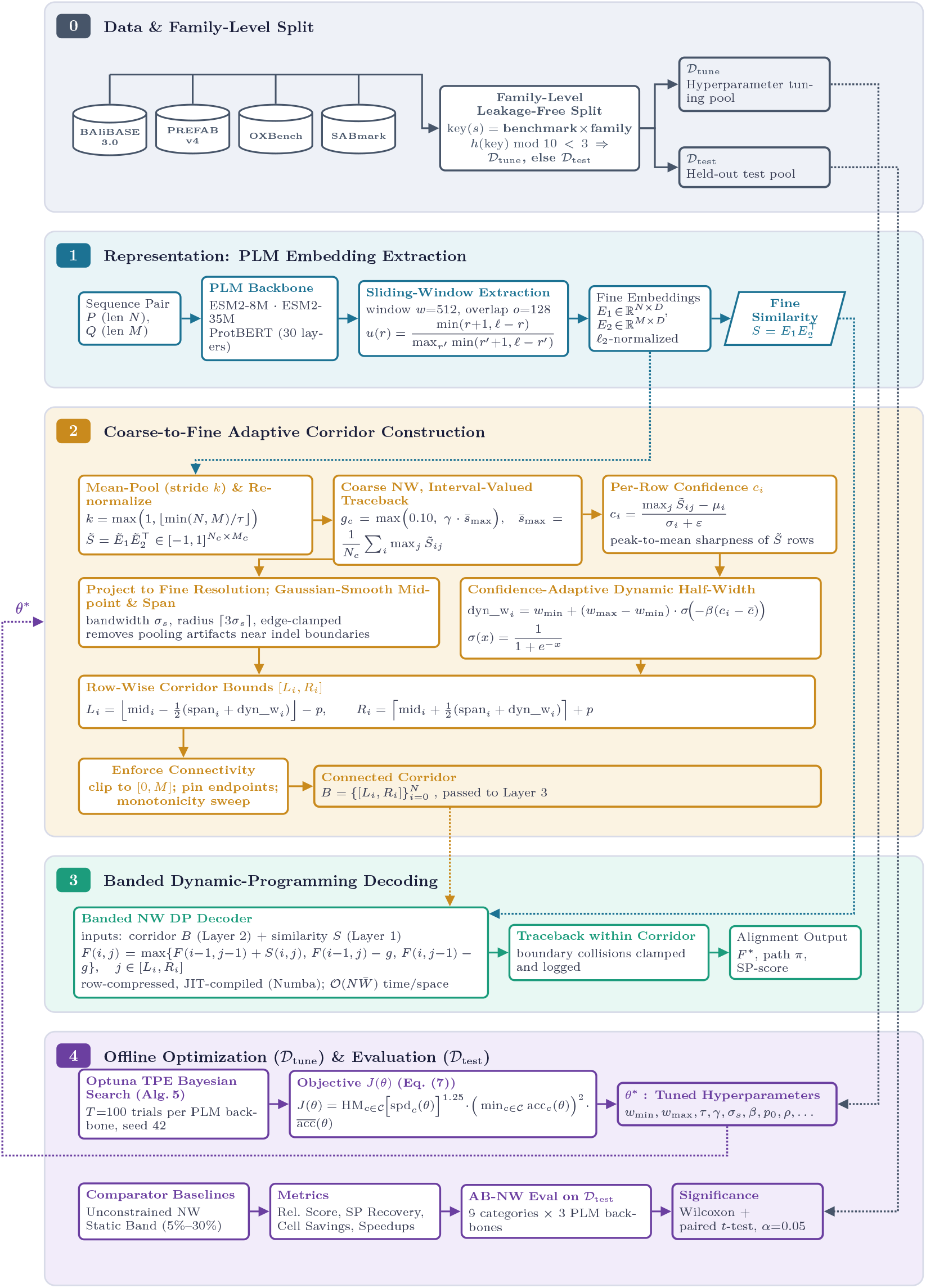
Architecture of AB-NW. **Layer 0:** Leakage-free family split (*D*_tune_ / *D*_test_). **Layer 1:** Sliding-window PLM embedding extraction. **Layer 2:** Coarse alignment and confidence-guided adaptive corridor construction ([*L*_*i*_, *R*_*i*_]). **Layer 3:** Row-compressed banded dynamic programming decoding. **Layer 4:** Offline Optuna TPE hyperparameter tuning and multi-category evaluation.

### 3.2. Protein Language Model Embedding Extraction

Residue embeddings derive from three pretrained PLM checkpoints: two ESM-2 transformer encoders (facebook/esm 2_t6_8M_UR50D, 6 layers; `facebook/esm2_t12_35M_UR50D`, 12 layers) [13], and one BERT-style model (`Rostlab/prot_b ert`, 30 layers) [14]. All models run in inference mode with mixed-precision autocast on CUDA. We discard boundary tokens from final hidden states and *ℓ*_2_-normalize each residue vector to yield *E* ∈ ℝ^*L*×*D*^.

To avoid *O*(*L*^2^) self-attention memory overhead on sequences exceeding *w* = 512, we process inputs in overlapping windows (*o* = 128, stride *s* = 384) and merge them via triangular weighting (Algorithm 1):

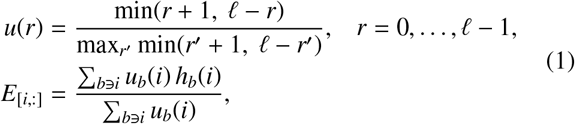

where *h*_*b*_(*i*) denotes the hidden state at position *i* in window *b*. Output rows are re-normalized to unit *ℓ*_2_ norm.

#### Algorithm 1 Sliding-Window Embedding Extraction

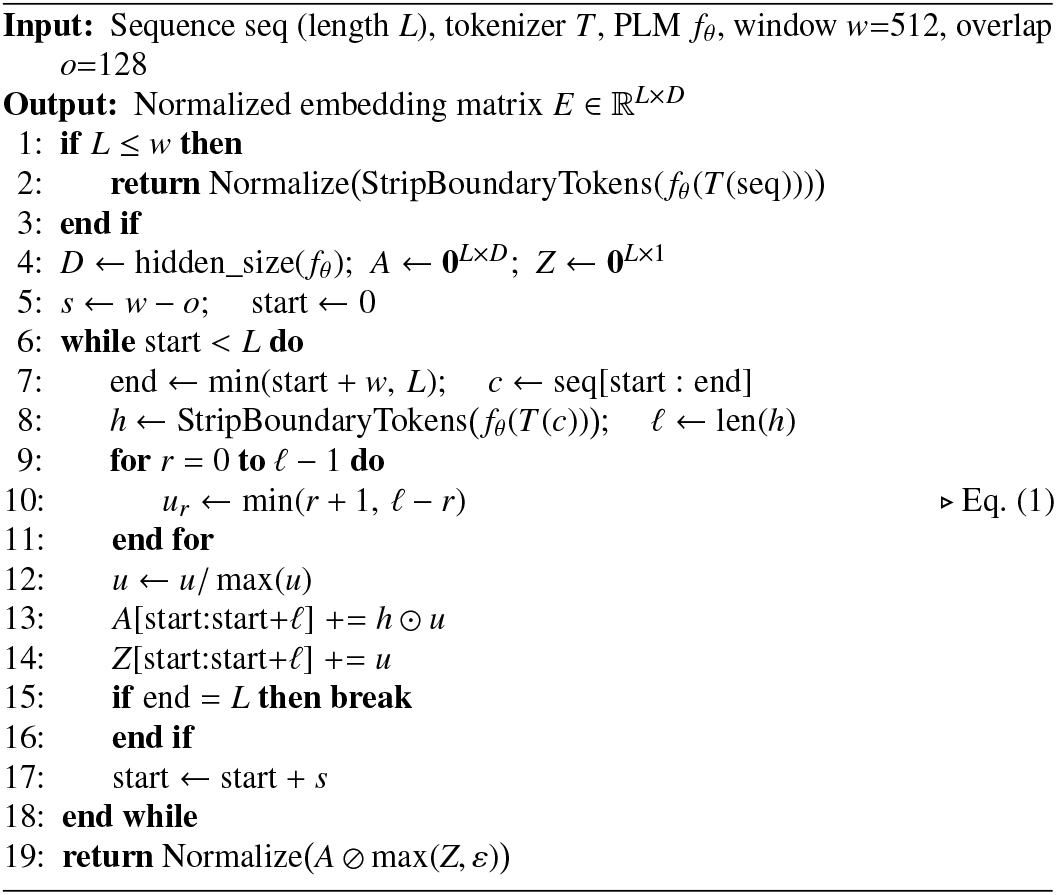

### 3.3. Coarse-to-Fine Adaptive Corridor Construction

#### 3.3.1. Coarse Global Alignment with Interval Traceback

Both embedding matrices are mean-pooled at stride *k* and *ℓ*_2_-normalized to yield 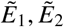, forming coarse cosine similarity 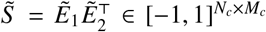 . We solve standard unbanded NW on 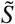 with linear gap penalty 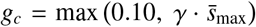, where 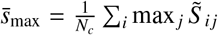. To retain near -tied candidate paths, the solver outputs a column interval 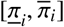 per row containing all columns touched during optimal traceback (Algorithm 2).

##### Algorithm 2 CorridorTracebackNW: Coarse Alignment

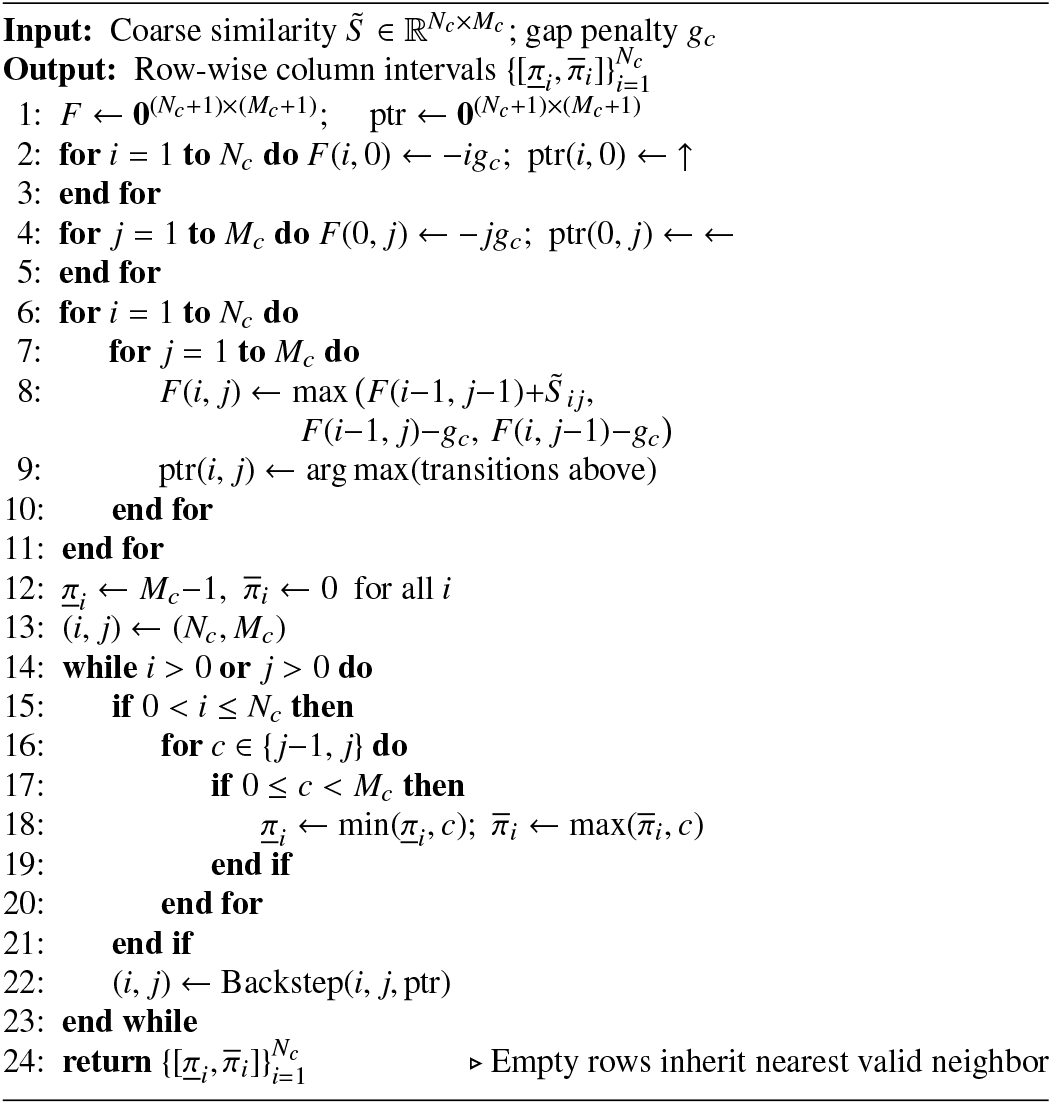

#### 3.3.2. Confidence-Adaptive Fine-Resolution Bounds

Projecting coarse intervals to fine resolution maps row *i* into *k* consecutive rows and scales column coordinates by *k*, with the final boundary mapped to *M*. Row confidence represents standardized peak-to-mean sharpness:

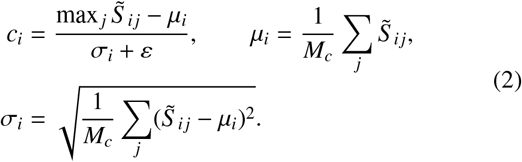

Diffuse rows yield low values; distinct matches yield high scores. Midpoint mid_*i*_ and span span_*i*_ undergo Gaussian smoothing (bandwidth *σ*_*s*_, support ⌈3*σ*_*s*_⌉, clamped boundaries) to remove pooling artifacts. Dynamic half-width interpolates between *w*_min_ and *w*_max_ based on confidence deviation:

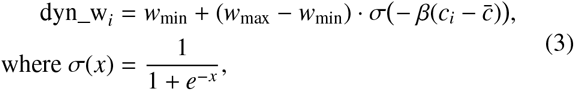

where *w*_max_ = clip max(*w*_max,min_, ⌊*Mκ*⌋), |*N* − *M*| + 2*p, M* , 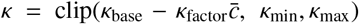, and safety padding *p* = *p*_0_ + ⌊*ρ* max(*N, M*)⌋ + *k*. Unconstrained bounds follow:

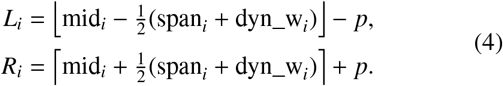

Bounds are clipped to [0, *M*], anchored at endpoints (*L*_0_=0, *R*_*N*_ =*M*), and filtered via forward–backward monotonicity sweeps to satisfy Definition 1 (Algorithm 3).

##### Algorithm 3 Coarse-to-Fine Adaptive Corridor Construction

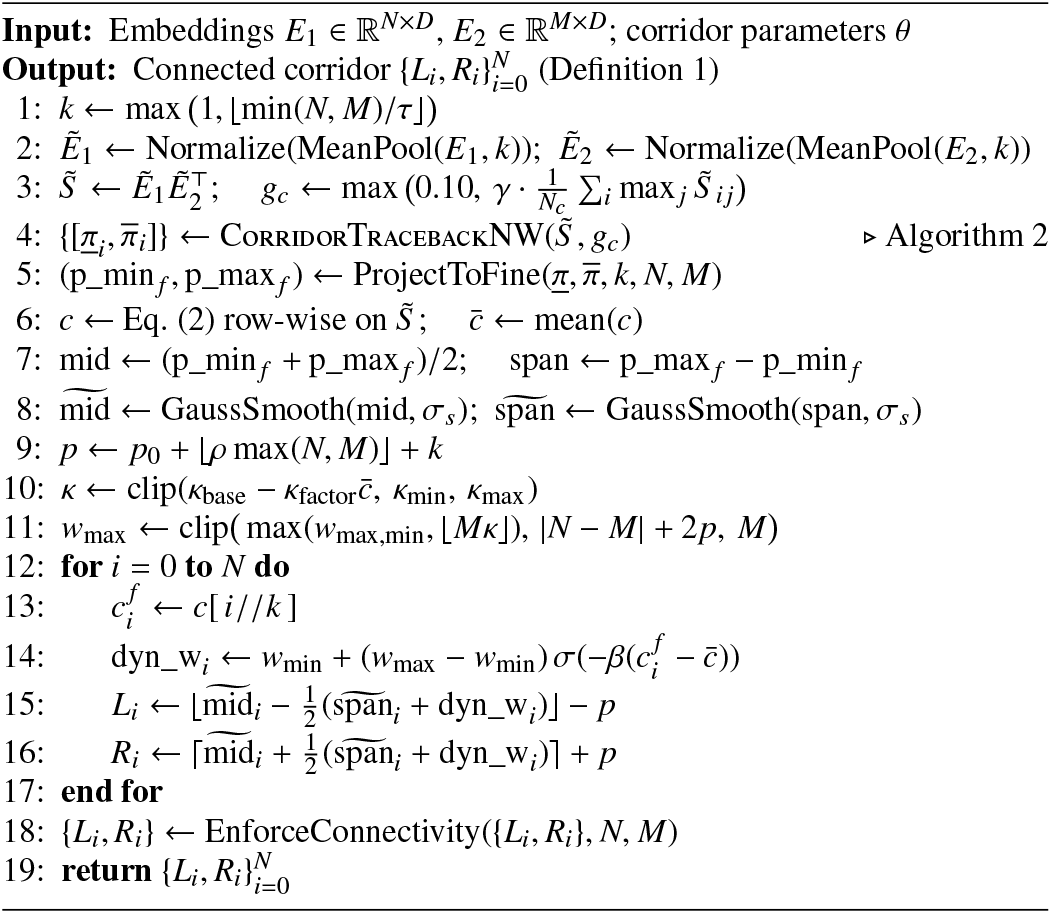

### 3.4. Banded Dynamic-Programming Decoder

Evaluating NW recursion across corridor *ℬ* operates on 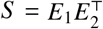 strictly within *j* ∈ [*L*_*i*_, *R*_*i*_]:

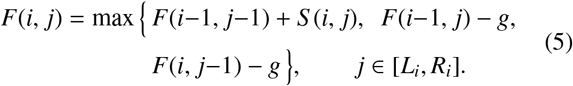

We set *g* = 0.15, with boundary values *F*(*i*, 0) = −*ig, F*(0, *j*) = − *jg*, and out-of-band transitions fixed to −∞. State values and pointers reside in row-compressed buffers of width *W*_max_ = max_*i*_ *w*_*i*_. If a traceback step exits *ℬ*, it clips to the nearest in-band boundary cell, ensuring continuous paths (Algorithm 4).

#### Algorithm 4 Banded Needleman–Wunsch Dynamic Programming

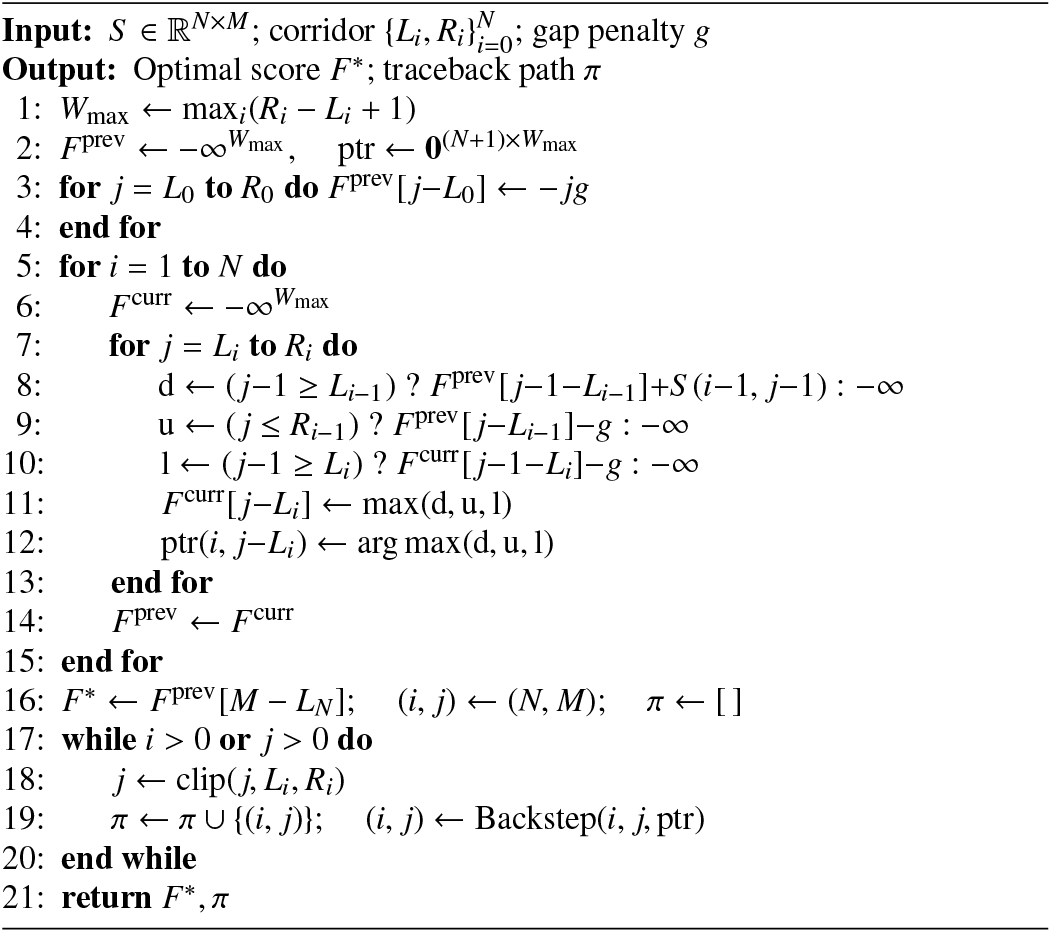

All pooling, smoothing, confidence scoring, boundary generation, and DP recursions are implemented in JIT-compiled Numba kernels (`fastmath`, `nogil`, row-parallelized). Kernels execute initial warmup cycles on dummy arrays to exclude JIT compilation overhead from recorded timings.

### 3.5. Generality of the Corridor to Other Alignment Recursions

Corridor construction depends strictly on embeddings *E*_1_, *E*_2_, operating independently of the cell-level dynamic programming recurrence. Replacing the global linear-gap recursion retains the *O*(*NW*_max_) visitation bound and row-compressed storage:

#### Local alignment (Smith–Waterman [23])

Setting *F*(*i, j*) = max{0, *F*(*i*−1, *j*−1) + *S* (*i, j*), *F*(*i*−1, *j*) − *g, F*(*i, j*−1) − *g*}, starting traceback at arg max_(*i, j*)∈*ℬ*_ *F*(*i, j*), and stopping at *F* = 0 bounds local alignment to *ℬ* in *O*(*NW*_max_) operations.

#### Affine gap penalties (Gotoh [10])

Replacing the single matrix with three coupled states for gap opening *o* and extension *e*,

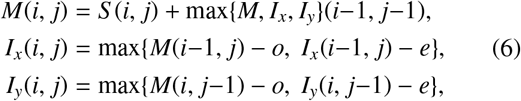

evaluated on *j* ∈ [*L*_*i*_, *R*_*i*_], yields *O*(3*NW*_max_) = *O*(*NW*_max_) execution.

Both variants retain Algorithms 1–3 and Proposition 2, requiring alterations only in lines 7–9 and 17 of Algorithm 4. We focus empirical benchmarks on global linear-gap NW (Eq. (5)), presenting these derivations as structural properties.

### 3.6. Optimality Guarantees and Complexity Analysis

#### Proposition 1

(Conditional Global Optimality). *Let* Π^*^(*P, Q*) *denote the set of globally optimal unconstrained alignment paths between sequences P and Q under the scoring system with score* 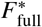. *If the connected corridor* 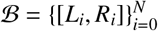 *contains at least one globally optimal path* 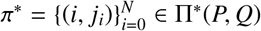 *such that L*_*i*_ ≤ *j*_*i*_ ≤ *R*_*i*_ *for all i* ∈ {0, … , *N*}, *then the banded dynamic programming decoder (Algorithm 4) computes* 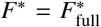 *and recovers an exact globally optimal alignment path π* ∈ Π^*^(*P, Q*).

*Proof*. Let *F*_full_(*i, j*) and *F*_*ℬ*_(*i, j*) denote dynamic programming values under unconstrained and banded decoding. We apply induction along the steps of *π*^*^.

*Base step:* Boundary condition *L*_0_ = 0 ≤ 0 ≤ *R*_0_ gives *F*_*ℬ*_(0, 0) = *F*_full_(0, 0) = 0. For all (0, *j*) ∈ *π*^*^, *F*_*ℬ*_(0, *j*) = − *jg* = *F*_full_(0, *j*).

*Inductive step:* Assume *F*_*ℬ*_(*i*^′^, *j*^′^) = *F*_full_(*i*^′^, *j*^′^) for predecessors (*i*^′^, *j*^′^) ∈ *π*^*^ prior to (*i, j*_*i*_). Because sub-paths of *π*^*^ remain optimal for sequence prefixes *P*_1:*i*_ and 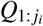:

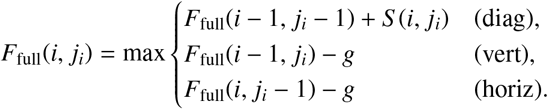

Because *π*^*^ ⊆ *ℬ*, the optimal predecessor satisfies 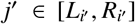. The induction hypothesis yields *F*_*ℬ*_(*i*^′^, *j*^′^) = *F*_full_(*i*^′^, *j*^′^). Algorithm 4 evaluates Eq. (5) across all valid in-band neighbors while routing out-of-band transitions to −∞, ensuring *F*_*ℬ*_(*i, j*_*i*_) ≥ *F*_full_(*i, j*_*i*_). Restricting search space cannot exceed the global optimum, giving *F*_*ℬ*_(*i, j*_*i*_) ≤ *F*_full_(*i, j*_*i*_). Thus, *F*_*ℬ*_(*i, j*_*i*_) = *F*_full_(*i, j*_*i*_).

By induction, 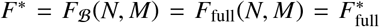 at the terminal cell (*N, M*). Backstepping along ptr recovers an optimal path *π* ∈ Π^*^(*P, Q*).

#### Proposition 2

(Time and Space Complexity of Banded Decoding). *For a connected corridor ℬ with mean row width* 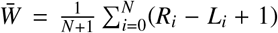 *and maximum row width W*_max_ = max_*i*_ *R*_*i*_ − *L*_*i*_ + 1), *Algorithm 4 executes in* 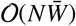 *time and requires* O(*NW*_max_) *space, compared to* O(*NM*) *time and space for the unconstrained baseline*.

*Proof*. The outer loop runs *N* times (line 3). In row *i*, the inner loop executes *R*_*i*_ − *L*_*i*_ + 1 iterations with *O*(1) operations per cell (line 5). The forward pass computes 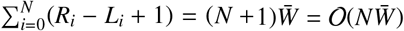 cell updates. Traceback inspects at most *N*+*M*+1 cells in *O*(1) steps each, remaining dominated by the forward sweep. Storing row-compressed pointers over (*N* + 1) × *W*_max_ cells bounds memory to O(*NW*_max_). Setting *L*_*i*_ = 0 and *R*_*i*_ = *M* for all *i* yields 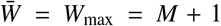, returning the classical unconstrained O(*NM*) baseline.

#### Corollary 1.

*Since W*_max_ ≤ *M by construction (Eq*. (4)*), banded DP is never asymptotically slower than unconstrained DP, achieving an algorithmic speedup factor bounded by* 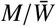, *maximized when confidence c*_*i*_ *(Eq*. (2)*) is uniformly high and dyn_w*_*i*_ → *w*_min_.

Coarse alignment dominates corridor construction, requiring *O*(*N*_*c*_ *M*_*c*_) = *O*(*τ*^2^) time. Because *τ* is constant, this term amortizes over the *O*(*N*) banded DP pass. Total pipeline complexity is 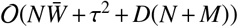 time and *O*(*NW*_max_) space, compared to *O*(*NM* + *D*(*N* + *M*)) time and space for unconstrained decoding.

#### 3.7. Hyperparameter Optimization

The corridor pipeline defines twelve hyperparameters *θ* (Table 2). We optimize *θ* separately per PLM backbone using Optuna’s Tree-structured Parzen Estimator (TPE) [59, 22] across *T* = 100 trials (seed 42). Each trial evaluates candidate configuration *θ*_*t*_ across all cached tuning pairs. Category accuracy acc_*c*_(*θ*_*t*_) and median speedup spd_*c*_(*θ*_*t*_) over classes C = {*c*_1_, … , *c*_9_} define the search objective:

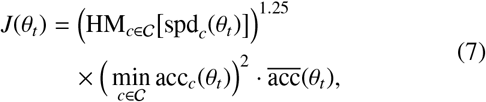

where 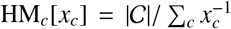 penalizes isolated slow categories, 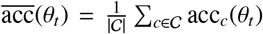 tracks mean accuracy, and *θ*^*^ = arg max_*t*_ *J*(*θ*_*t*_). The 1.25 exponent prioritizes narrow bands; the squared minimum accuracy term prevents degradation on difficult subsets (Algorithm 5).

Table 3 lists parameter bounds explored during tuning. Table 4 reports selected values *θ*^*^.

**Table 3:** Optuna search space for corridor hyperparameters.

| Parameter (Alias) | Search Range | Step |
| --- | --- | --- |
| $w_{\min}$ ($w_{\min}$) | [2, 8] (int) | 2 |
| $w_{\max, \min}$ ($w_{\max, \min}$) | [6, 16] (int)<sup>a</sup> | 2 |
| $\kappa_{\min}$ ($\text{conf\_scale\_min}$) | [0.01, 0.05] | 0.01 |
| $\kappa_{\max}$ ($\text{conf\_scale\_max}$) | [0.04, 0.10]<sup>b</sup> | 0.01 |
| $\tau$ ($\text{target\_coarse\_len}$) | [35, 65] (int) | 5 |
| $\gamma$ ($\text{coarse\_gap\_factor}$) | [0.10, 0.30] | 0.05 |
| $\sigma_s$ ($\text{smooth\_sigma}$) | [1.0, 3.0] | 0.5 |
| $p_0$ ($\text{padding}$) | [0, 3] (int) | 1 |
| $\rho$ ($\text{adapt\_pad\_scale}$) | [0.0001, 0.0008] | 0.0001 |
| $\kappa_{\text{base}}$ ($\text{conf\_scale\_base}$) | [0.05, 0.12] | 0.01 |
| $\kappa_{\text{factor}}$ ($\text{conf\_scale\_factor}$) | [0.01, 0.03] | 0.01 |
| $\beta$ ($\text{conf\_sigmoid\_beta}$) | [1.2, 2.5] | 0.1 |

<sup>a</sup>Floored at $w_{\min} + 2$ . <sup>b</sup>Floored at $\kappa_{\min} + 0.01$ .

**Table 4:** Per-model tuned corridor hyperparameters *θ*^*^ selected by Algorithm 5 (*T* =100 trials, seed 42). Gap penalty *g* = 0.15.

| Hyperparameter | ESM2-8M | ESM2-35M | ProtBERT |
| --- | --- | --- | --- |
| $w_{\min}$ | 6 | 8 | 8 |
| $w_{\max, \min}$ | 12 | 14 | 12 |
| $\kappa_{\min}$ | 0.01 | 0.03 | 0.01 |
| $\kappa_{\max}$ | 0.08 | 0.07 | 0.05 |
| $\tau$ | 65 | 45 | 40 |
| $\gamma$ | 0.10 | 0.10 | 0.30 |
| $\sigma_s$ | 3.0 | 1.0 | 3.0 |
| $p_0$ | 3 | 1 | 3 |
| $\rho$ | 0.0008 | 0.0007 | 0.0001 |
| $\kappa_{\text{base}}$ | 0.07 | 0.11 | 0.10 |
| $\kappa_{\text{factor}}$ | 0.03 | 0.03 | 0.01 |
| $\beta$ | 2.2 | 2.4 | 1.2 |
| Best objective $J(\theta^*)$ | 0.796 | 0.497 | 0.453 |

##### Algorithm 5

Bayesian Optimization of Corridor Parameters

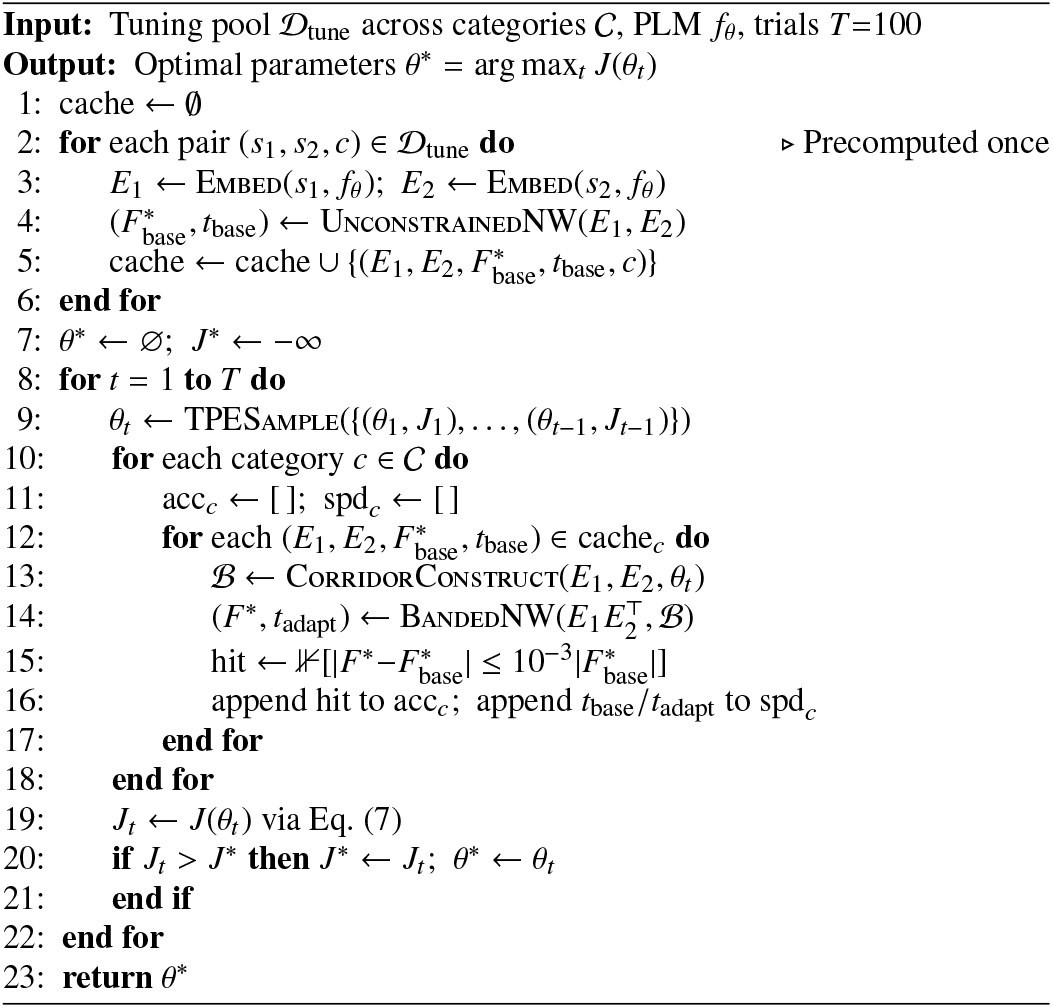

### 3.8. Reference Alignment Corpus and Data-Leakage Control

Reference benchmarks derive from the Edgar repository (drive5.com/bench), which unifies BAliBASE v3 [18], PRE-FAB v4 [19], OXBench [20], and SABmark [21]. We project each MSA onto pairwise alignments, remove double-gap columns, and preserve Sum-of-Pairs core masks [60, 19] (*m* ∈ {0, 1}^|aln|^, marked by upper-case residues). Sequence pairs satisfy 1 ≤ *N, M* ≤ 20 000 and core coverage ∑*m* ≥ 5. Three metrics characterize alignment difficulty:

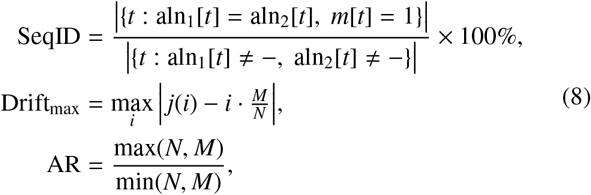

where *j*(*i*) indexes the reference column aligned with residue *i*.

Pairs are organized into nine benchmark categories (Table 5):

**Table 5:** Stratified benchmark evaluation corpus (D_test_, 400 sequence pairs across 9 categories) derived via family-level leakage-free split from BAliBASE, SABmark, PREFAB, and OXBench. Full candidate pool sizes and metric distributions are provided in Supplementary Tables S9 and S10.

| Category | Selection Criteria | Pairs | Mean Dimensions ($N \times M$) | SeqID (%) | Drift (res) | Aspect Ratio |
| --- | --- | --- | --- | --- | --- | --- |
| 1. Twilight Zone | SeqID $\leq 30\%$, Drift<sub>max</sub> $\geq 20$ | 50 | 528 × 559 | 20.3% ± 3.9% | 77.2 ± 71.3 | 1.35 ± 0.42 |
| 2. Ultra-Giant | $N \times M \geq 3 \times 10^5$, max size | 25 | 8481 × 8481 | 47.9% ± 43.5% | 0.0 ± 0.0 | 1.00 ± 0.00 |
| 3. Diverse Large | $N \times M \geq 3 \times 10^5$, family-capped | 25 | 3795 × 3795 | 49.1% ± 29.4% | 0.0 ± 0.0 | 1.00 ± 0.00 |
| 4. Asymmetric Indels | $\vert N - M\vert \geq 100$ residues | 50 | 321 × 343 | 20.5% ± 16.5% | 119.8 ± 68.8 | 2.41 ± 0.74 |
| 5. Extreme Ratio | Aspect Ratio $\geq 2.5$ | 50 | 203 × 244 | 21.4% ± 19.1% | 99.5 ± 89.5 | 3.08 ± 0.59 |
| 6. Internal Repeats | Tandem repeats / 4-mer count $\geq 8$ | 50 | 806 × 806 | 54.7% ± 17.7% | 0.0 ± 0.0 | 1.00 ± 0.00 |
| 7. Micro-Peptides | min($N, M$) $\leq 80$ residues | 50 | 62 × 62 | 55.7% ± 31.2% | 1.2 ± 3.0 | 1.03 ± 0.08 |
| 8. Standard Bench | Both lengths in [200, 1000] | 50 | 538 × 538 | 53.3% ± 17.9% | 0.0 ± 0.0 | 1.00 ± 0.00 |
| 9. Unbiased Control | Uniform random sampling | 50 | 708 × 709 | 52.4% ± 20.2% | 1.0 ± 4.9 | 1.01 ± 0.05 |

#### Matrix-size prioritization (Categories 1–3)

Twilight Zone, Ultra-Giant, and Diverse Large Matrix pools are ranked by matrix area *N* × *M*. Top-area pairs are selected, with Category 3 applying a family-level frequency cap to enforce structural variety.

#### Targeted feature filtering (Categories 4–8)

Specific thresholds isolate asymmetric indels, extreme aspect ratios, internal tandem repeats, micro-peptides, and standard bench-mark dimensions.

#### Unbiased control (Category 9)

Uniform random sampling across reference alignments provides a control baseline.

##### Definition 2

(Family-level leakage-free split). Let key(*s*) = `benchmark_name-alignment_set_name` denote the source MSA family for pair *s*, and *h*(·) the low-order 32 bits of its MD5 hash. Pair *s* enters the tuning pool (*D*_tune_) if *h*(key(*s*)) mod 10 *<* 3; otherwise, it joins test set (*D*_test_).

Partitioning strictly by MSA identifier eliminates cross-split data leakage: no test pair shares a homologous family with the tuning set. Table 5 details the evaluation split (*D*_test_). Supplementary Tables S9 and S10 provide candidate pool breakdowns and source distributions.

### 3.9. Comparator Methods

We compare three alignment approaches on identical PLM embeddings:

1. **Unconstrained baseline**. Standard global NW over unconstrained matrix 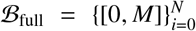, establishing ground-truth score 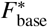.
2. **Static band**. Uniform diagonal corridor *L*_*i*_ = max(0, ⌊*iM/N*⌋ − *w/*2), *R*_*i*_ = min(*M*, ⌊*iM/N*⌋ + *w/*2), evaluated at *w* = max(10, ⌊*M f* ⌋) for *f* ∈ {5%, 10%, 15%, 20%, 25%, 30%} [30].
3. **Adaptive band (proposed)**. Dynamic corridor *ℬ* (Algorithm 3) with backbone-specific parameters *θ*^*^ (Algorithm 5, Table 4).

All methods share identical DP kernels (Algorithm 4), isolating corridor geometry as the sole experimental variable.

### 3.10. Implementation and Hardware Environment

Experiments ran on Linux 6.12 (glibc 2.35) with Python 3.12.13, PyTorch 2.10.0 (CUDA 12.8) [61], Hugging Face `transformers` 5.0.0 [62], `optuna` 4.9.0, and `numba` 0.60.0 (`fastmath`, `nogil`) [63]. PLM inference ran across dual NVIDIA Tesla T4 GPUs (16 GB VRAM each); corridor construction and DP decoding executed on the host CPU. A global random seed of 42 fixed all stochastic components.

## 4. Results and Discussion

We evaluate AB-NW against the unconstrained O(*NM*) global DP baseline and six static-banding configurations (*w* ∈ {5%, 10%, 15%, 20%, 25%, 30%}) across three PLMs: ESM2-8M (`facebook/esm2_t6_8M_UR50D`), ESM2-35M (`facebo ok/esm2_t12_35M_UR50D`), and ProtBERT (`Rostlab/prot_bert`).

Sequence lengths span 60 to 8,500 residues. Because pooling across divergent sequence topologies skews aggregate metrics toward long pairs, all evaluations, cell counts, wall-clock measurements, and hypothesis tests are reported strictly per category and backbone. Table 6 outlines cross-category performance. Table 7 isolates Category 4 (Asymmetric Indels) to demonstrate static-banding failure. Supplementary Tables S1 to S8 provide complete comparator breakdowns for the remaining categories.

**Table 6:** Cross-category performance summary of AB-NW across nine challenge regimes and three PLM backbones. Values denote Mean ± Standard Deviation across held-out test pairs (D_test_). Supplementary Tables S1 to S8 provide individual static banding comparisons (5% to 30%).

| Benchmark Challenge Category | PLM Backbone | Rel Score (%) | Rel SP (%) | DP Cell Savings (%) | Pure DP Speedup | End-to-End Speedup |
| --- | --- | --- | --- | --- | --- | --- |
| **1. Twilight Zone**<br>($N = 50$, SeqID $\leq 30\%$, Drift $\geq 20$) | ESM-35M | 99.94 | 99.43 | $77.5\% \pm 10.8\%$ | $6.23 \times \pm 1.90\times$ | $1.39 \times \pm 0.44\times$ |
| | ESM-8M | 99.93 | 99.91 | $78.8\% \pm 11.9\%$ | $6.21 \times \pm 2.97\times$ | $1.44 \times \pm 0.43\times$ |
| | ProtBERT | 99.98 | 100.00 | $76.3\% \pm 8.8\%$ | $7.63 \times \pm 3.67\times$ | $1.32 \times \pm 0.66\times$ |
| **2. Ultra-Giant Matrices**<br>($N = 25$, avg. $8481 \times 8481$) | ESM-35M | 100.00 | 100.00 | $88.3\% \pm 1.1\%$ | $9.79 \times \pm 0.95\times$ | $2.27 \times \pm 0.10\times$ |
| | ESM-8M | 100.00 | 100.00 | $91.6\% \pm 0.8\%$ | $12.60 \times \pm 1.22\times$ | $2.89 \times \pm 0.13\times$ |
| | ProtBERT | 100.00 | 100.00 | $87.7\% \pm 0.1\%$ | $11.67 \times \pm 0.36\times$ | $1.73 \times \pm 0.05\times$ |
| **3. Diverse Large Matrices**<br>($N = 25$, family-capped, avg. $3795 \times 3795$) | ESM-35M | 99.22 | 90.56 | $87.9\% \pm 0.8\%$ | $10.10 \times \pm 0.75\times$ | $2.35 \times \pm 0.13\times$ |
| | ESM-8M | 99.91 | 91.75 | $91.7\% \pm 0.5\%$ | $13.30 \times \pm 0.93\times$ | $2.94 \times \pm 0.15\times$ |
| | ProtBERT | 100.00 | 100.00 | $87.6\% \pm 0.1\%$ | $12.89 \times \pm 2.72\times$ | $1.82 \times \pm 0.51\times$ |
| **4. Asymmetric Indels**<br>($N = 50$, $\vert N - M\vert = 243 \pm 90$) | ESM-35M | 99.61 | 98.99 | $60.2\% \pm 8.8\%$ | $7.66 \times \pm 13.61\times$ | $1.40 \times \pm 1.55\times$ |
| | ESM-8M | 99.87 | 99.88 | $60.9\% \pm 8.8\%$ | $5.74 \times \pm 7.16\times$ | $1.21 \times \pm 1.12\times$ |
| | ProtBERT | 99.81 | 105.37 | $59.1\% \pm 10.9\%$ | $6.49 \times \pm 8.14\times$ | $1.02 \times \pm 0.94\times$ |
| **5. Extreme Ratio Asymmetry**<br>($N = 50$, Aspect Ratio $\geq 2.5$) | ESM-35M | 99.75 | 99.94 | $58.2\% \pm 7.3\%$ | $11.15 \times \pm 19.34\times$ | $1.42 \times \pm 2.13\times$ |
| | ESM-8M | 99.80 | 99.26 | $57.7\% \pm 7.6\%$ | $8.02 \times \pm 11.97\times$ | $1.16 \times \pm 1.55\times$ |
| | ProtBERT | 99.81 | 99.52 | $55.3\% \pm 10.2\%$ | $8.01 \times \pm 9.45\times$ | $0.89 \times \pm 1.01\times$ |
| **6. Internal Repeats**<br>($N = 50$, avg. $806 \times 806$) | ESM-35M | 100.00 | 100.00 | $86.7\% \pm 1.6\%$ | $11.03 \times \pm 8.80\times$ | $1.67 \times \pm 1.07\times$ |
| | ESM-8M | 100.00 | 100.00 | $89.6\% \pm 2.2\%$ | $11.73 \times \pm 6.34\times$ | $1.86 \times \pm 0.93\times$ |
| | ProtBERT | 100.00 | 100.00 | $85.2\% \pm 2.1\%$ | $10.79 \times \pm 5.18\times$ | $1.10 \times \pm 0.60\times$ |
| **7. Short Micro-Peptides**<br>($N = 50$, $\min(N, M) \leq 80$) | ESM-35M | 100.00 | 100.00 | $70.1\% \pm 5.8\%$ | $55.58 \times \pm 117.22\times$ | $2.54 \times \pm 5.26\times$ |
| | ESM-8M | 100.00 | 100.00 | $67.0\% \pm 6.6\%$ | $16.14 \times \pm 27.59\times$ | $0.84 \times \pm 1.39\times$ |
| | ProtBERT | 100.00 | 100.00 | $65.7\% \pm 6.9\%$ | $34.88 \times \pm 66.68\times$ | $1.50 \times \pm 3.02\times$ |
| **8. Standard Benchmark Suite**<br>($N = 50$, avg. $538 \times 538$) | ESM-35M | 100.00 | 100.00 | $86.9\% \pm 0.6\%$ | $11.26 \times \pm 8.29\times$ | $1.69 \times \pm 1.14\times$ |
| | ESM-8M | 100.00 | 100.00 | $89.8\% \pm 1.3\%$ | $10.13 \times \pm 3.52\times$ | $1.60 \times \pm 0.77\times$ |
| | ProtBERT | 100.00 | 100.00 | $85.2\% \pm 1.4\%$ | $11.73 \times \pm 11.15\times$ | $1.17 \times \pm 0.87\times$ |
| **9. Unbiased Random Control**<br>($N = 50$, avg. $708 \times 709$) | ESM-35M | 100.00 | 100.00 | $85.7\% \pm 3.8\%$ | $18.12 \times \pm 33.26\times$ | $2.05 \times \pm 2.20\times$ |
| | ESM-8M | 100.00 | 100.00 | $88.6\% \pm 5.0\%$ | $12.80 \times \pm 13.65\times$ | $1.79 \times \pm 1.04\times$ |
| | ProtBERT | 100.00 | 100.00 | $84.1\% \pm 4.7\%$ | $13.52 \times \pm 17.74\times$ | $1.22 \times \pm 1.02\times$ |

**Table 7:** Exemplar comparator breakdown on Category 4: Asymmetric Indels (*N* = 50 pairs, |*N* − *M*| = 243 ± 90), showing static banding breakdown versus AB-NW. Supplementary Tables S1 to S8 report other categories. Values represent Mean ± Standard Deviation.

| Model | Method | E2E Spd. | DP Spd. | Cell Sav.% | Rel Score% | Rel SP% | SP-Score |
| --- | --- | --- | --- | --- | --- | --- | --- |
| **ESM-35M** | Unconstrained | 1.00x | 1.00x | 0.0% | 100.00 | 100.00 | $0.6218 \pm 0.3827$ |
| | Static (5%) | $4.23x \pm 4.95x$ | $29.03x \pm 42.75x$ | $94.3\% \pm 1.5\%$ | 82.46 | 29.19 | $0.1815 \pm 0.3463$ |
| | Static (10%) | $4.07x \pm 4.85x$ | $20.46x \pm 31.52x$ | $90.2\% \pm 0.3\%$ | 86.18 | 40.72 | $0.2532 \pm 0.3648$ |
| | Static (15%) | $3.74x \pm 4.40x$ | $14.73x \pm 21.72x$ | $85.6\% \pm 0.2\%$ | 88.89 | 48.07 | $0.2989 \pm 0.3831$ |
| | Static (20%) | $3.44x \pm 4.13x$ | $11.83x \pm 17.41x$ | $81.0\% \pm 0.2\%$ | 90.75 | 56.48 | $0.3512 \pm 0.3819$ |
| | Static (25%) | $3.35x \pm 4.29x$ | $10.66x \pm 17.11x$ | $76.6\% \pm 0.2\%$ | 92.40 | 61.27 | $0.3810 \pm 0.3835$ |
| | Static (30%) | $3.11x \pm 3.87x$ | $9.32x \pm 14.72x$ | $72.2\% \pm 0.2\%$ | 93.95 | 66.95 | $0.4163 \pm 0.3786$ |
|  | **AB-NW (Ours)** | **$1.40x \pm 1.55x$** | **$7.66x \pm 13.61x$** | **$60.2\% \pm 8.8\%$** | **99.61** | **98.99** | **$0.6155 \pm 0.3899$** |
| **ESM-8M** | Unconstrained | 1.00x | 1.00x | 0.0% | 100.00 | 100.00 | $0.4902 \pm 0.4280$ |
| | Static (5%) | $3.96x \pm 3.65x$ | $21.82x \pm 24.90x$ | $94.3\% \pm 1.5\%$ | 85.48 | 35.45 | $0.1738 \pm 0.3457$ |
| | Static (10%) | $3.73x \pm 3.67x$ | $15.86x \pm 20.97x$ | $90.2\% \pm 0.3\%$ | 88.27 | 48.25 | $0.2365 \pm 0.3697$ |
| | Static (15%) | $3.29x \pm 3.18x$ | $12.00x \pm 16.15x$ | $85.6\% \pm 0.2\%$ | 90.51 | 55.04 | $0.2698 \pm 0.3891$ |
| | Static (20%) | $3.29x \pm 3.56x$ | $9.67x \pm 12.23x$ | $81.0\% \pm 0.2\%$ | 91.96 | 64.42 | $0.3158 \pm 0.3890$ |
| | Static (25%) | $2.98x \pm 2.98x$ | $8.48x \pm 11.94x$ | $76.6\% \pm 0.2\%$ | 92.99 | 70.58 | $0.3460 \pm 0.4019$ |
| | Static (30%) | $2.88x \pm 3.01x$ | $7.51x \pm 10.16x$ | $72.2\% \pm 0.2\%$ | 93.94 | 74.05 | $0.3630 \pm 0.4038$ |
|  | **AB-NW (Ours)** | **$1.21x \pm 1.12x$** | **$5.74x \pm 7.16x$** | **$60.9\% \pm 8.8\%$** | **99.87** | **99.88** | **$0.4896 \pm 0.4309$** |
| **ProtBERT** | Unconstrained | 1.00x | 1.00x | 0.0% | 100.00 | 100.00 | $0.3428 \pm 0.4338$ |
| | Static (5%) | $2.54x \pm 2.23x$ | $25.31x \pm 21.47x$ | $94.3\% \pm 1.5\%$ | 89.59 | 34.68 | $0.1189 \pm 0.2995$ |
| | Static (10%) | $2.52x \pm 2.09x$ | $18.07x \pm 20.02x$ | $90.2\% \pm 0.3\%$ | 92.00 | 43.96 | $0.1507 \pm 0.3231$ |
| | Static (15%) | $2.32x \pm 2.06x$ | $12.97x \pm 13.03x$ | $85.6\% \pm 0.2\%$ | 93.49 | 47.35 | $0.1623 \pm 0.3225$ |
| | Static (20%) | $2.25x \pm 1.82x$ | $10.78x \pm 11.71x$ | $81.0\% \pm 0.2\%$ | 94.36 | 57.73 | $0.1979 \pm 0.3428$ |
| | Static (25%) | $2.16x \pm 1.91x$ | $9.17x \pm 11.77x$ | $76.6\% \pm 0.2\%$ | 95.17 | 59.48 | $0.2039 \pm 0.3458$ |
| | Static (30%) | $2.10x \pm 1.63x$ | $8.24x \pm 10.29x$ | $72.2\% \pm 0.2\%$ | 95.88 | 64.64 | $0.2216 \pm 0.3536$ |
|  | **AB-NW (Ours)** | **$1.02x \pm 0.94x$** | **$6.49x \pm 8.14x$** | **$59.1\% \pm 10.9\%$** | **99.81** | **105.37** | **$0.3612 \pm 0.4413$** |

Reported metrics comprise:

i. **Relative DP Score Ratio (%):** 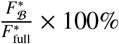.
ii. **Cell Savings (%):** 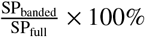.
iii. **Relative SP Score Recovery (%):** 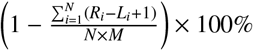.
iv. **Speedup Ratios:** Pure DP Speedup (*t*_base,DP_*/t*_banded,DP_) and End-to-End Speedup (*t*_base,total_*/t*_adapt,total_), where the latter includes pooling, confidence estimation, smoothing, and corridor generation.

### 4.1. Category-Specific Comparative Performance Analysis

#### 4.1.1. Low Identity and Off-Diagonal Drift Regimes (Categories 1, 4, and 5)

Categories 1, 4, and 5 expose the structural failure points of static diagonal bands.

Category 1 (Twilight Zone) pairs exhibit low identity (mean 20.3% ± 3.9%) and large off-diagonal drift (mean 77.2 ± 71.3 residues). Static corridors centered on *j* = *i*· *M/N* clip these drifting paths (Supplementary Figure S1B), recovering only 50.71% SP core blocks under ESM-35M (Supplementary Table S1) and degrading alignment score from 223.4 to 194.7 on `1oyc_2tmdA`. AB-NW widens dynamically via interval coarse traceback (Supplementary Figure S1C, D). It recovers 99.43% (ESM-35M), 99.91% (ESM-8M), and 100.00% (ProtBERT) SP core blocks, retaining 99.93% to 99.98% of the unconstrained DP score while discarding 76.3% to 78.8% of cells (*p <* 0.05; Supplementary Table S11).

Category 4 (Asymmetric Indels) pairs contain substantial insertions or deletions (|*N* −*M*| = 243±90 residues). Static bands bisect indel loops (Figure 2B), dropping the `BB20028_p1_5` score from 223.8 to 214.3. Across Category 4, static 5% banding recovers only 29.19% SP, and static 30% banding reaches only 66.95% SP under ESM-35M (Table 7). AB-NW routes search corridors around these loops (Figure 2C, D), achieving 98.99% SP recovery on ESM-35M, 99.88% on ESM-8M, and 105.37% on ProtBERT (Table 7). The *>* 100% ProtBERT recovery occurs because semantic embeddings guide the corridor toward true structural equivalences that standard unconstrained linear-gap DP penalizes incorrectly.

**Figure 2:**
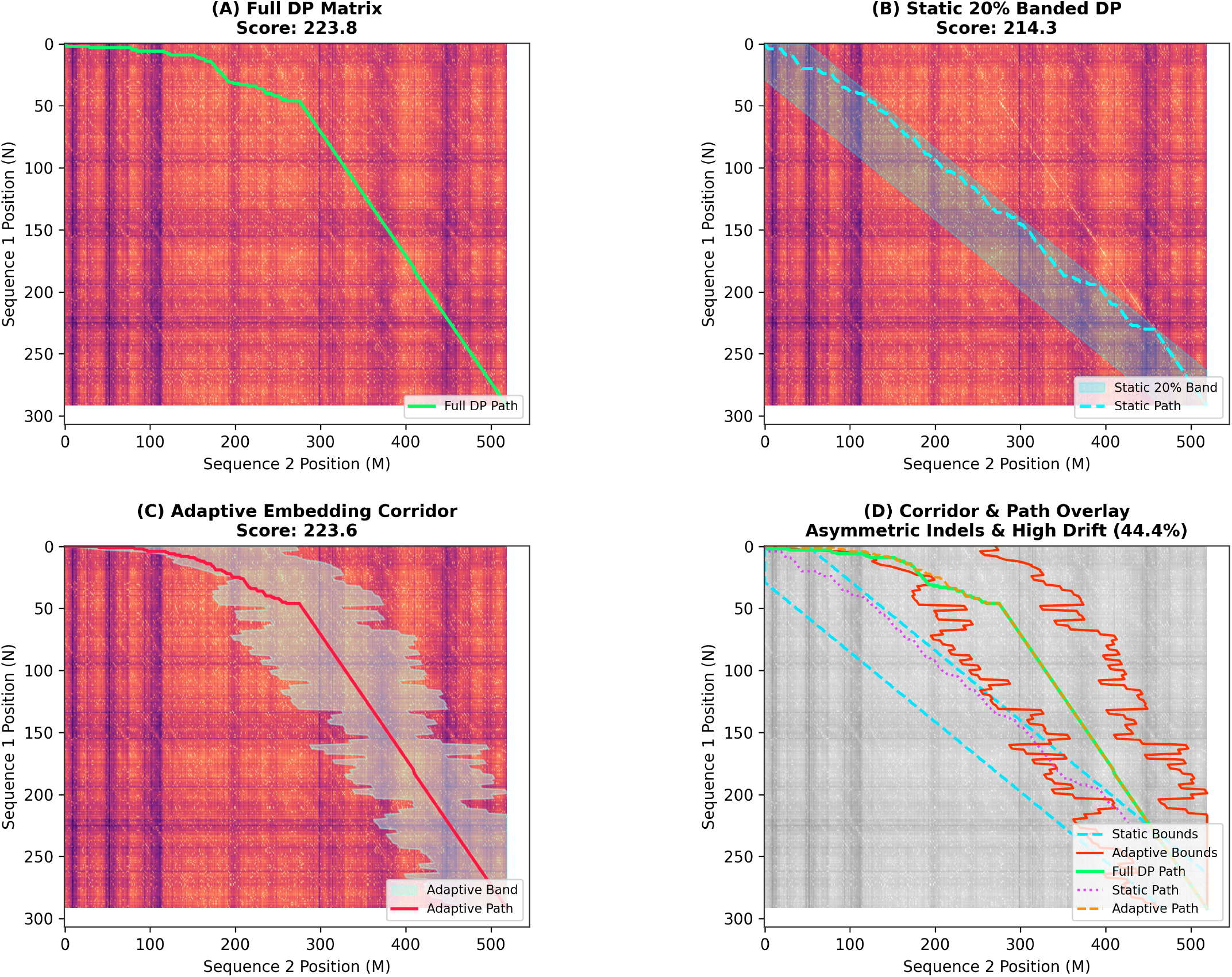
Alignment search on pair BB20028_p1_5 exhibiting asymmetric indels and 44.4% drift (300 × 520 matrix) from Category 4. **(A)** Unconstrained DP matrix (score: 223.8). **(B)** Static 20% band bisects insertion loops, dropping score to 214.3. **(C)** Adaptive corridor widens around insertions, attaining a 223.6 score. **(D)** Overlay of static versus adaptive search corridors.

Category 5 (Extreme Ratio Asymmetry) features steep aspect ratios (mean AR = 3.08 ± 0.59). Static bands truncate alignment paths along skewed slopes (Supplementary Figure S2B), yielding only 20.39% to 25.05% SP recovery at 5% width (Supplementary Table S4). AB-NW expands row width *w*_*i*_ adaptively (Supplementary Figure S2C, D), matching the unconstrained score of 47.6 on pair 1krs_1lylA and restoring 99.26% to 99.94% SP recovery across backbones (Table 6) with 8.01× to 11.15× Pure DP speedups. ProtBERT End-to-End speedup drops to 0.89× here because 30-layer inference overhead dominates runtime on these smaller matrices (203 × 244 average; Section 4.4).

#### 4.1.2. Large Matrix Scale and Asymptotic Acceleration Regimes (Categories 2 and 3)

Categories 2 and 3 evaluate performance on large matrices (*N* × *M* ≥ 3 × 10^5^ cells).

Category 2 (Ultra-Giant Matrices) scales to 8, 481 × 8, 481 residues (7.19 × 10^7^ cells), where unconstrained DP consumes 1.02 to 1.44 s per pair. High embedding confidence across long sequences yields compact corridors. AB-NW achieves 100.00% score and SP recovery across all backbones (Table 6; Supplementary Table S2). Speedups reach 12.60× Pure DP (2.89× End-to-End) on ESM-8M and 11.67× Pure DP on ProtBERT, reducing execution time from 1.4390 s to 0.8302 s by pruning 87.7% to 91.6% of cells (*p <* 10^−7^; Supplementary Table S11).

Category 3 (Diverse Large Matrices) caps representation at two pairs per family across 3, 795 × 3, 795 average dimensions. AB-NW yields 13.30× Pure DP speedup (2.94× End-to-End) on ESM-8M and 12.89× Pure DP on ProtBERT, retaining *>* 99.2% score fidelity (Table 6; Supplementary Table S3). Under ESM-35M, AB-NW SP recovery is 90.56% whereas static bands score 104.40% across widths. Because static bands fully enclose these specific trajectories, they inherit the reference alignment topology directly; AB-NW tightens bounds to optimize the exact DP score (99.22% relative) rather than structural SP agreement.

#### 4.1.3. Symmetric, Repetitive, and Micro-Peptide Regimes (Categories 6, 7, 8, and 9)

Category 6 (Internal Repeats) evaluates tandem duplications (806 × 806 average size). Static bands and AB-NW both attain 100.00% score recovery (Supplementary Table S5). AB-NW ignores off-diagonal suboptimal repeat matches, eliminating 85.2% to 89.6% of cells to deliver 10.79× to 11.73× Pure DP speedups (Table 6).

Category 7 (Short Micro-Peptides) averages 62 × 62 residues (*<* 3, 844 cells, *<* 0.1 ms DP runtime). Model forward passes and tensor memory transfers dominate runtime. End-to-End speedups span 0.84× to 2.54×, despite Pure DP speedups reaching 16.14× to 55.58× (Table 6; Supplementary Table S6). Coarse boundary rounding degrades static band recovery to 96.72% SP, whereas AB-NW achieves 100.00% score and SP recovery.

Categories 8 (Standard Benchmark Suite) and 9 (Unbiased Random Control) represent typical lengths (538 × 538 and 708 × 709 residues). AB-NW provides 10.13× to 18.12× Pure DP speedups (1.17× to 2.05× End-to-End) and 100.00% score and SP recovery across all backbones (Table 6; Supplementary Tables S7 and S8).

### 4.2 Synthesizing the Trade-off: When Does Adaptive Banding vs. Static Banding Excel?

Empirical results delineate boundary conditions governing static versus adaptive heuristics:

a. **Where Static Banding Suffers:** Linear projections (*j* = *i* · *M/N*) fail whenever sequences exhibit low sequence identity with non-linear drift (Category 1), asymmetric indel loops with |*N* − *M*| ≥ 100 (Category 4), or severe length skew with AR ≥ 2.5 (Category 5). Static 5% to 10% corridors clip optimal paths in these domains, dropping SP score recovery to between 20% and 50% (Table 7; Supplementary Tables S1 and S4).
b. **Where Static Banding is Acceptable:** Highly homologous, length-matched sequence pairs with strictly diagonal trajectories (Categories 6, 8, and 9) execute accurately within static 15% to 20% bands, avoiding dynamic corridor setup costs (Supplementary Tables S5, S7, and S8).
c. **The Blind Real-World Selection Dilemma and AB-NW’s Advantage:** In uncurated, large-scale workflows such as metagenomic clustering, structural properties like drift, indels, and aspect ratios remain unknown *a priori*. Choosing narrow static bands (5% to 10%) risks path clipping and alignment collapse; choosing wide bands (25% to 30%) foregoes practical acceleration.
d. **The Adaptive Solution:** AB-NW removes this manual tuning requirement. It contracts search widths 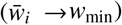 across confident diagonal alignments to accelerate decoding, while automatically widening across indel-heavy or drifting pairs to preserve optimal alignment scores.

### 4.3. Cross-Model Comparison and Embedding Topology Effects

AB-NW operates across model architectures, though its performance reflects the underlying representation geometry:

- **Model Scale and Representation Capacity:** ESM2-35M (12 layers, *D* = 480) generates sharper similarity matrices than ESM2-8M (6 layers, *D* = 320). On Category 2, ESM2-35M achieves higher mean confidence (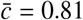 versus 0.74 for ESM2-8M), producing tighter corridors (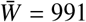 versus 1085 residues) and 88.3% cell savings compared to 87.7% on ProtBERT (Table 6).
- **ProtBERT vs. ESM Representation Topologies:** Prot-BERT (30 layers, *D* = 1024) outputs diffuse similarity fields. Bayesian HPO compensates by selecting a larger coarse gap factor (*γ* = 0.30 for ProtBERT versus *γ* = 0.10 for ESM backbones; Table 4), which sharpens the coarse alignment.
- **Semantic Alignment Overriding Linear Gap Penalties:** In Category 4, ProtBERT with AB-NW yields a 105.37% relative SP recovery relative to unconstrained linear-gap DP (Table 7). Contextual embeddings identify structural correspondences across large loops that fixed affine penalties misalign.

### 4.4. Computational Amortization and Profiling Bottlenecks

Profiled runtimes partition into two scaling regimes:

- **Small Sequence Regime** (*N* ≤ 100): In Category 7, DP execution requires *<* 0.1 ms. PyTorch model evaluation and tensor conversions dominate total runtime, capping End-to-End speedups at 0.84× to 2.54× despite 16.14× to 55.58× pure DP acceleration (Table 6; Supplementary Table S6).
- **Asymptotic Amortization Regime** (*N* ≥ 1, 000): Quadratic DP costs dominate Category 2 and 3 runtimes. Coarse pooling overhead (*O*(*τ*^2^) with *τ* ∈ [40, 65]) drops below 0.5% of runtime, yielding 2.27× to 2.94× End-to-End speedups and 10× to 13.3× pure DP reductions (Table 6).
- **Memory Footprint Savings:** Row-compressed allocation with width *W*_max_ = max_*i*_ *w*_*i*_ reduces memory demand from O(*NM*) to *O*(*NW*_max_), executing 20, 000 × 20, 000 global alignments without memory faults.
- **ProtBERT Cache Utilization:** Naive cell models predict speedups of 1*/*(1 − savings). ESM backbones adhere to this relation, but ProtBERT exceeds it; in Category 3, 87.6% cell savings predicts ∼ 8.1× versus 12.89× observed, and in Category 8, 85.2% predicts ∼ 6.8× versus 11.73× observed. ProtBERT’s diffuse scores and *γ* = 0.30 parameter produce uniform corridor profiles, enhancing CPU cache locality.

### 4.5. Hyperparameter Sensitivity and HPO Convergence

Optuna Bayesian optimization (Algorithm 5, *T* = 100 trials, seed 42) converged to backbone-specific configurations *θ*^*^ (Table 4):

- **Target Coarse Stride** *τ*: Tuned to *τ* = 65 for ESM2-8M, *τ* = 45 for ESM2-35M, and *τ* = 40 for ProtBERT. Finer sampling resolves diffuse ProtBERT representations; ESM models use coarse strides, lowering the O(*τ*^2^) search cost.
- **Sigmoid Sharpness** *β*: Selected at *β* = 2.4 for ESM2-35M, *β* = 2.2 for ESM2-8M, and *β* = 1.2 for ProtBERT. Steeper sigmoid slopes contract confident regions toward *w*_min_ more aggressively.
- **Safety Padding (***p*_0_, *ρ*): Base offsets *p*_0_ ∈ [1, 3] and scaling factors *ρ* ∈ [0.0001, 0.0008] prevent corridor boundary collisions during projection.

### 4.6. Statistical Significance and Hypothesis Testing

Statistical validation comprised 54 paired tests across the nine benchmark categories and three PLM backbones, using Pratt-modified Wilcoxon signed-rank tests and paired *t*-tests at *α* = 0.05.

#### 4.6.1. Confirmation of Quality Hypothesis 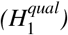

AB-NW outperforms static 20% banding across all off-diagonal test categories (Supplementary Table S11):

- **Category 4 (Asymmetric Indels):** AB-NW increases SP core recovery over static 20% bands by +26.43% on ESM2-35M (*p* = 3.68 × 10^−7^, *d* = 0.8374), +17.38% on ESM2-8M (*p* = 0.00048, *d* = 0.6118), and +16.33% on ProtBERT (*p* = 0.00045, *d* = 0.5182).
- **Category 5 (Extreme Ratio Asymmetry):** AB-NW gains +27.76% SP score on ESM2-35M (*p* = 3.07 × 10^−7^, *d* = 0.9072), +17.65% on ESM2-8M (*p* = 5.91 × 10^−5^, *d* = 0.6385), and +13.46% on ProtBERT (*p* = 0.00015, *d* = 0.5055).
- **Category 1 (Twilight Zone):** AB-NW shows significant SP improvements under low sequence identity (+10.47% for ESM2-35M, *p* = 0.0055, *d* = 0.3975; +3.29% for ProtBERT, *p* = 0.0254, *d* = 0.2413).

#### 4.6.2. Confirmation of Speedup Hypothesis 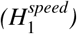

Across all nine categories and three backbones, pure DP execution time drops significantly below the unconstrained baseline (*p <* 10^−3^ to *p <* 10^−15^, *W* = 0.0; Supplementary Table S11). Category 2 shows large absolute execution drops, saving −1.0090 s per pair on ESM2-35M (*p* = 5.96 × 10^−8^, *d* = −25.2742) and −1.3157 s on ProtBERT (*d* = −31.1170).

## 5. Conclusion

In this study, we proposed the AB-NW framework, which unifies deep contextual language representations with exact DP. By decoupling search-space restriction from the downstream DP scoring function, AB-NW resolves a long-standing trade-off in biological sequence alignment: the choice between computationally expensive unconstrained DP (O(*NM*)) and fragile static banding heuristics (O(*NW*)) that fail when alignment paths deviate from the diagonal trajectory. By leveraging coarse-to-fine embedding projections and standardized confidence signals *c*_*i*_ prior to full-resolution decoding, the corridor dynamically contracts in high-confidence regions 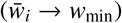 while expanding and recentering across ambiguous or indel-rich regions to safely enclose the optimal path.

This formulation demonstrates that pruning the dynamic programming search space does not require sacrificing alignment fidelity or exact optimization guarantees. As established theoretically, banded decoding over a connected corridor *ℬ* guarantees exact DP global optimality whenever *ℬ* encloses the true optimal trajectory (Proposition 1). In benchmark categories where static 5%–20% bands suffer catastrophic alignment collapse—recovering only 20%–50% of core-block SP scores in the Twilight Zone, asymmetric indels, and extreme aspect ratios—AB-NW consistently recovers *>* 98.9% of reference SP accuracy while eliminating 55.3%–78.8% of active DP cells across PLM backbones. Furthermore, because coarse-alignment overhead amortizes to less than 0.5% on large sequences (*N, M* ≥ 3,700), storing value and pointer matrices in row-compressed buffers achieves an effective 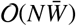 runtime and O(*NW*_max_) memory footprint (Proposition 2), yielding end-to-end speedups of up to 2.94× (13.30× in pure DP) and enabling global alignment of giant protein sequences (*>* 8,000 residues) on standard hardware.

Importantly, the corridor-construction mechanism depends solely on embedding similarities (*E*_1_, *E*_2_) rather than the DP scoring recurrence itself. As a result, the row-compressed enumeration grid is mathematically compatible with local alignment (Smith–Waterman) and multi-state affine-gap recurrences (Gotoh) without altering the O(*NW*_max_) visitation bound. While end-to-end throughput on short micro-peptides (min(*N, M*) ≤ 80) remains constrained by PyTorch forward passes and tensor transfer latencies, future work will address this by deploying quantized PLM encoders and compiling row-compressed DP kernels directly onto GPU wavefront architectures. Ultimately, AB-NW bridges the gap between deep contextual representations and constrained DP, making exact, sequence-adaptive alignment practical for large-scale, high-throughput pipelines.

## Data and Source Code Availability Statement

The reference alignment benchmark collections parsed in this study are publicly available from the Edgar benchmark repository (http://drive5.com/bench). The source code and execution scripts implementing the AB-NW algorithm and benchmark pipeline are publicly available on GitHub at https://github.com/Shoaib-CS/Adaptive-Banding-Needleman-Wunsch-PLM.

## Declaration of generative AI and AI-assisted technologies in the manuscript preparation process

During the preparation of this work, the authors used Google Gemini in order to improve the language throughout the manuscript. After using this tool, the authors reviewed and edited the content as needed and take full responsibility for the content of the published article.

## Ethics Approval and Consent to Participate

Not applicable. This study is a computational analysis based on publicly available datasets and did not involve human participants or animal subjects.

## Consent for Publication

Not applicable.

