## Supplementary Material for "Pruning the Search, Not the Signal: Adaptive-Banding Needleman–Wunsch Sequence Alignment via Protein Language Model Confidence"

**Abstract**

This document provides supplementary benchmark tables, database provenance breakdowns, candidate selection statistics, and complete 7-method comparator evaluations supporting the main manuscript.

### S1. Supplementary Benchmark Results: Full Baseline Comparisons

The following tables and figures provide full comparator evaluations, dataset breakdowns, statistical tests, and visual alignment corridors supporting the main manuscript:

- **Table S1:** Category 1: Twilight Zone
- **Table S2:** Category 2: Ultra-Giant Matrices
- **Table S3:** Category 3: Diverse Large Matrices
- **Table S4:** Category 5: Extreme Ratio Asymmetry
- **Table S5:** Category 6: Internal Repeats
- **Table S6:** Category 7: Short Micro-Peptides
- **Table S7:** Category 8: Standard Benchmark Suite
- **Table S8:** Category 9: Unbiased Random Control
- **Table S9:** Stratified Benchmark Category Candidate Pool and Sampled Splits ($\mathcal{D}_{\text{tune}}$ and $\mathcal{D}_{\text{test}}$)
- **Table S10:** Aggregated Structural Characteristics, Coarse Downsampling Stride ($k$), and Database Provenance
- **Table S11:** Paired Statistical Hypothesis Tests Across Backbones and Categories
- **Figure S1:** Visual Alignment Corridor on Twilight Zone Drift Pair (`1oyc_2tmdA`)
- **Figure S2:** Visual Alignment Corridor on Extreme Aspect Ratio Pair (`1krs_1lylA`)

*Corresponding author.

Table S1: Empirical alignment metrics on **Category 1: Twilight Zone** ($N$=50 pairs, average matrix size 528 × 559). Values reported as Mean ± Standard Deviation.

| Model | Method | E2E Spd. | DP Spd. | Cell Sav.% | Rel Score% | Rel SP% | SP-Score |
|---|---|---|---|---|---|---|---|
| **ESM-35M** | Unconstrained | 1.00x | 1.00x | 0.0% | 100.00 | 100.00 | 0.7011 ± 0.3237 |
| | Static (5%) | 2.91x ± 1.21x | 16.49x ± 6.42x | 95.0% ± 0.1% | 83.51 | 50.71 | 0.3555 ± 0.3856 |
| | Static (10%) | 2.70x ± 1.05x | 10.35x ± 3.63x | 90.2% ± 0.1% | 90.54 | 67.76 | 0.4751 ± 0.3980 |
| | Static (15%) | 2.48x ± 0.99x | 7.98x ± 2.85x | 85.5% ± 0.1% | 94.44 | 79.15 | 0.5549 ± 0.3715 |
| | Static (20%) | 2.35x ± 0.90x | 6.61x ± 2.34x | 81.0% ± 0.1% | 96.38 | 84.48 | 0.5923 ± 0.3604 |
| | Static (25%) | 2.22x ± 0.83x | 5.56x ± 1.93x | 76.6% ± 0.1% | 97.39 | 87.62 | 0.6143 ± 0.3505 |
| | Static (30%) | 2.08x ± 0.79x | 4.85x ± 1.70x | 72.2% ± 0.1% | 98.15 | 90.84 | 0.6369 ± 0.3326 |
| | **AB-NW (Ours)** | **1.39x ± 0.44x** | **6.23x ± 1.90x** | **77.5% ± 10.8%** | **99.94** | **99.43** | **0.6971 ± 0.3290** |
| **ESM-8M** | Unconstrained | 1.00x | 1.00x | 0.0% | 100.00 | 100.00 | 0.6647 ± 0.3354 |
| | Static (5%) | 3.46x ± 0.79x | 15.32x ± 3.57x | 95.0% ± 0.1% | 89.28 | 61.56 | 0.4092 ± 0.3955 |
| | Static (10%) | 2.91x ± 0.70x | 8.95x ± 1.85x | 90.2% ± 0.1% | 94.40 | 80.76 | 0.5368 ± 0.4044 |
| | Static (15%) | 2.70x ± 0.65x | 6.94x ± 1.69x | 85.5% ± 0.1% | 97.56 | 93.91 | 0.6242 ± 0.3520 |
| | Static (20%) | 2.52x ± 0.65x | 5.73x ± 1.46x | 81.0% ± 0.1% | 99.06 | 96.89 | 0.6440 ± 0.3464 |
| | Static (25%) | 2.31x ± 0.55x | 4.89x ± 1.21x | 76.6% ± 0.1% | 99.41 | 97.46 | 0.6478 ± 0.3473 |
| | Static (30%) | 2.13x ± 0.50x | 4.20x ± 0.93x | 72.2% ± 0.1% | 99.58 | 97.46 | 0.6478 ± 0.3473 |
| | **AB-NW (Ours)** | **1.44x ± 0.43x** | **6.21x ± 2.97x** | **78.8% ± 11.9%** | **99.93** | **99.91** | **0.6641 ± 0.3312** |
| **ProtBERT** | Unconstrained | 1.00x | 1.00x | 0.0% | 100.00 | 100.00 | 0.7191 ± 0.3290 |
| | Static (5%) | 2.60x ± 1.29x | 25.67x ± 14.78x | 95.0% ± 0.1% | 94.16 | 57.78 | 0.4155 ± 0.3963 |
| | Static (10%) | 2.49x ± 1.30x | 14.72x ± 7.28x | 90.2% ± 0.1% | 97.96 | 81.23 | 0.5841 ± 0.4176 |
| | Static (15%) | 2.34x ± 1.22x | 10.81x ± 5.11x | 85.5% ± 0.1% | 99.28 | 92.05 | 0.6619 ± 0.3626 |
| | Static (20%) | 2.16x ± 1.11x | 8.82x ± 4.27x | 81.0% ± 0.1% | 99.84 | 95.44 | 0.6863 ± 0.3573 |
| | Static (25%) | 2.07x ± 1.18x | 7.31x ± 3.85x | 76.6% ± 0.1% | 99.97 | 96.19 | 0.6917 ± 0.3585 |
| | Static (30%) | 2.02x ± 1.03x | 6.45x ± 3.10x | 72.2% ± 0.1% | 99.97 | 96.19 | 0.6917 ± 0.3585 |
| | **AB-NW (Ours)** | **1.32x ± 0.66x** | **7.63x ± 3.67x** | **76.3% ± 8.8%** | **99.98** | **100.00** | **0.7191 ± 0.3290** |

Table S2: Empirical alignment metrics on **Category 2: Ultra-Giant Matrices** ($N$=25 pairs, average matrix size 8481 × 8481). Values reported as Mean ± Standard Deviation.

| Model | Method | E2E Spd. | DP Spd. | Cell Sav.% | Rel Score% | Rel SP% | SP-Score |
|---|---|---|---|---|---|---|---|
| **ESM-35M** | Unconstrained | 1.00x | 1.00x | 0.0% | 100.00 | 100.00 | 0.9759 ± 0.0182 |
| | Static (5%) | 2.86x ± 0.15x | 22.63x ± 1.38x | 95.1% ± 0.0% | 100.00 | 100.00 | 0.9759 ± 0.0182 |
| | Static (10%) | 2.53x ± 0.14x | 11.74x ± 0.51x | 90.2% ± 0.0% | 100.00 | 100.00 | 0.9759 ± 0.0182 |
| | Static (15%) | 2.36x ± 0.08x | 8.15x ± 0.29x | 85.6% ± 0.0% | 100.00 | 100.00 | 0.9759 ± 0.0182 |
| | Static (20%) | 2.20x ± 0.06x | 6.57x ± 0.23x | 81.0% ± 0.0% | 100.00 | 100.00 | 0.9759 ± 0.0182 |
| | Static (25%) | 2.06x ± 0.10x | 5.54x ± 0.23x | 76.6% ± 0.0% | 100.00 | 100.00 | 0.9759 ± 0.0182 |
| | Static (30%) | 1.96x ± 0.07x | 4.88x ± 0.13x | 72.2% ± 0.0% | 100.00 | 100.00 | 0.9759 ± 0.0182 |
| | **AB-NW (Ours)** | **2.27x ± 0.10x** | **9.79x ± 0.95x** | **88.3% ± 1.1%** | **100.00** | **100.00** | **0.9759 ± 0.0182** |
| **ESM-8M** | Unconstrained | 1.00x | 1.00x | 0.0% | 100.00 | 100.00 | 0.9768 ± 0.0236 |
| | Static (5%) | 3.43x ± 0.10x | 20.93x ± 0.65x | 95.1% ± 0.0% | 100.00 | 100.00 | 0.9768 ± 0.0236 |
| | Static (10%) | 2.99x ± 0.08x | 10.84x ± 0.31x | 90.2% ± 0.0% | 100.00 | 100.00 | 0.9768 ± 0.0236 |
| | Static (15%) | 2.63x ± 0.10x | 7.42x ± 0.26x | 85.6% ± 0.0% | 100.00 | 100.00 | 0.9768 ± 0.0236 |
| | Static (20%) | 2.45x ± 0.06x | 6.06x ± 0.10x | 81.0% ± 0.0% | 100.00 | 100.00 | 0.9768 ± 0.0236 |
| | Static (25%) | 2.29x ± 0.05x | 5.11x ± 0.08x | 76.6% ± 0.0% | 100.00 | 100.00 | 0.9768 ± 0.0236 |
| | Static (30%) | 2.13x ± 0.06x | 4.45x ± 0.09x | 72.2% ± 0.0% | 100.00 | 100.00 | 0.9768 ± 0.0236 |
| | **AB-NW (Ours)** | **2.89x ± 0.13x** | **12.60x ± 1.22x** | **91.6% ± 0.8%** | **100.00** | **100.00** | **0.9768 ± 0.0236** |
| **ProtBERT** | Unconstrained | 1.00x | 1.00x | 0.0% | 100.00 | 100.00 | 0.9806 ± 0.0263 |
| | Static (5%) | 2.05x ± 0.07x | 28.19x ± 1.19x | 95.1% ± 0.0% | 100.00 | 100.00 | 0.9806 ± 0.0263 |
| | Static (10%) | 1.94x ± 0.06x | 14.58x ± 0.60x | 90.2% ± 0.0% | 100.00 | 100.00 | 0.9806 ± 0.0263 |
| | Static (15%) | 1.82x ± 0.07x | 10.17x ± 0.29x | 85.6% ± 0.0% | 100.00 | 100.00 | 0.9806 ± 0.0263 |
| | Static (20%) | 1.76x ± 0.07x | 8.32x ± 0.21x | 81.0% ± 0.0% | 100.00 | 100.00 | 0.9806 ± 0.0263 |
| | Static (25%) | 1.70x ± 0.05x | 7.02x ± 0.23x | 76.6% ± 0.0% | 100.00 | 100.00 | 0.9806 ± 0.0263 |
| | Static (30%) | 1.64x ± 0.04x | 6.09x ± 0.14x | 72.2% ± 0.0% | 100.00 | 100.00 | 0.9806 ± 0.0263 |
| | **AB-NW (Ours)** | **1.73x ± 0.05x** | **11.67x ± 0.36x** | **87.7% ± 0.1%** | **100.00** | **100.00** | **0.9806 ± 0.0263** |

Table S3: Empirical alignment metrics on **Category 3: Diverse Large Matrices** ($N$=25 pairs, average matrix size $3795 \times 3795$). Values reported as Mean ± Standard Deviation.

| Model | Method | E2E Spd. | DP Spd. | Cell Sav.% | Rel Score% | Rel SP% | SP-Score |
|---|---|---|---|---|---|---|---|
| **ESM-35M** | Unconstrained | 1.00x | 1.00x | 0.0% | 100.00 | 100.00 | 0.8473 ± 0.2667 |
| | Static (5%) | 3.19x ± 0.33x | 22.85x ± 1.34x | 95.1% ± 0.0% | 99.31 | 104.40 | 0.8846 ± 0.2002 |
| | Static (10%) | 2.84x ± 0.28x | 12.46x ± 1.10x | 90.2% ± 0.0% | 99.31 | 104.40 | 0.8846 ± 0.2002 |
| | Static (15%) | 2.56x ± 0.29x | 8.56x ± 0.79x | 85.6% ± 0.0% | 99.31 | 104.40 | 0.8846 ± 0.2002 |
| | Static (20%) | 2.39x ± 0.17x | 6.64x ± 0.57x | 81.0% ± 0.0% | 99.31 | 104.40 | 0.8846 ± 0.2002 |
| | Static (25%) | 2.22x ± 0.17x | 5.48x ± 0.45x | 76.6% ± 0.0% | 99.31 | 104.40 | 0.8846 ± 0.2002 |
| | Static (30%) | 2.06x ± 0.16x | 4.62x ± 0.36x | 72.2% ± 0.0% | 99.31 | 104.40 | 0.8846 ± 0.2002 |
| | **AB-NW (Ours)** | **2.35x ± 0.13x** | **10.10x ± 0.75x** | **87.9% ± 0.8%** | **99.22** | **90.56** | **0.7673 ± 0.3498** |
| **ESM-8M** | Unconstrained | 1.00x | 1.00x | 0.0% | 100.00 | 100.00 | 0.9694 ± 0.0910 |
| | Static (5%) | 3.82x ± 0.38x | 20.84x ± 1.80x | 95.1% ± 0.0% | 100.00 | 100.00 | 0.9694 ± 0.0910 |
| | Static (10%) | 3.23x ± 0.38x | 11.60x ± 0.76x | 90.2% ± 0.0% | 100.00 | 100.00 | 0.9694 ± 0.0910 |
| | Static (15%) | 2.93x ± 0.21x | 7.97x ± 0.57x | 85.6% ± 0.0% | 100.00 | 100.00 | 0.9694 ± 0.0910 |
| | Static (20%) | 2.63x ± 0.20x | 6.08x ± 0.49x | 81.0% ± 0.0% | 100.00 | 100.00 | 0.9694 ± 0.0910 |
| | Static (25%) | 2.38x ± 0.22x | 4.96x ± 0.43x | 76.6% ± 0.0% | 100.00 | 100.00 | 0.9694 ± 0.0910 |
| | Static (30%) | 2.19x ± 0.12x | 4.22x ± 0.29x | 72.2% ± 0.0% | 100.00 | 100.00 | 0.9694 ± 0.0910 |
| | **AB-NW (Ours)** | **2.94x ± 0.15x** | **13.30x ± 0.93x** | **91.7% ± 0.5%** | **99.91** | **91.75** | **0.8894 ± 0.2826** |
| **ProtBERT** | Unconstrained | 1.00x | 1.00x | 0.0% | 100.00 | 100.00 | 0.9980 ± 0.0102 |
| | Static (5%) | 2.30x ± 0.29x | 28.93x ± 3.81x | 95.1% ± 0.0% | 100.00 | 100.00 | 0.9980 ± 0.0102 |
| | Static (10%) | 2.15x ± 0.31x | 16.04x ± 3.63x | 90.2% ± 0.0% | 100.00 | 100.00 | 0.9980 ± 0.0102 |
| | Static (15%) | 1.99x ± 0.19x | 10.81x ± 1.46x | 85.6% ± 0.0% | 100.00 | 100.00 | 0.9980 ± 0.0102 |
| | Static (20%) | 1.87x ± 0.17x | 8.27x ± 0.64x | 81.0% ± 0.0% | 100.00 | 100.00 | 0.9980 ± 0.0102 |
| | Static (25%) | 1.90x ± 0.52x | 7.08x ± 1.53x | 76.6% ± 0.0% | 100.00 | 100.00 | 0.9980 ± 0.0102 |
| | Static (30%) | 1.83x ± 0.55x | 6.04x ± 1.34x | 72.2% ± 0.0% | 100.00 | 100.00 | 0.9980 ± 0.0102 |
| | **AB-NW (Ours)** | **1.82x ± 0.51x** | **12.89x ± 2.72x** | **87.6% ± 0.1%** | **100.00** | **100.00** | **0.9980 ± 0.0102** |

Table S4: Empirical alignment metrics on **Category 5: Extreme Ratio Asymmetry** ($N$=50 pairs, aspect ratio AR $\geq$ 2.5). Values reported as Mean ± Standard Deviation.

| Model | Method | E2E Spd. | DP Spd. | Cell Sav.% | Rel Score% | Rel SP% | SP-Score |
|---|---|---|---|---|---|---|---|
| **ESM-35M** | Unconstrained | 1.00x | 1.00x | 0.0% | 100.00 | 100.00 | 0.5277 ± 0.3819 |
| | Static (5%) | 4.44x ± 6.54x | 33.89x ± 52.83x | 93.2% ± 3.2% | 78.48 | 21.38 | 0.1128 ± 0.2082 |
| | Static (10%) | 4.41x ± 6.37x | 27.19x ± 44.10x | 89.7% ± 1.7% | 83.11 | 33.09 | 0.1746 ± 0.2501 |
| | Static (15%) | 4.27x ± 6.22x | 22.29x ± 37.12x | 85.4% ± 0.8% | 86.18 | 41.29 | 0.2179 ± 0.3037 |
| | Static (20%) | 3.64x ± 5.51x | 17.74x ± 28.94x | 80.9% ± 0.3% | 88.92 | 47.34 | 0.2498 ± 0.3157 |
| | Static (25%) | 3.96x ± 5.94x | 15.79x ± 25.61x | 76.6% ± 0.3% | 90.74 | 56.24 | 0.2968 ± 0.3336 |
| | Static (30%) | 3.87x ± 5.91x | 14.07x ± 23.62x | 72.3% ± 0.4% | 92.01 | 63.65 | 0.3359 ± 0.3513 |
| | **AB-NW (Ours)** | **1.42x ± 2.13x** | **11.15x ± 19.34x** | **58.2% ± 7.3%** | **99.75** | **99.94** | **0.5274 ± 0.3816** |
| **ESM-8M** | Unconstrained | 1.00x | 1.00x | 0.0% | 100.00 | 100.00 | 0.4195 ± 0.3866 |
| | Static (5%) | 3.91x ± 4.92x | 24.47x ± 31.83x | 93.2% ± 3.2% | 79.55 | 25.05 | 0.1051 ± 0.2085 |
| | Static (10%) | 3.83x ± 5.25x | 19.42x ± 25.26x | 89.7% ± 1.7% | 83.52 | 38.62 | 0.1620 ± 0.2563 |
| | Static (15%) | 3.68x ± 5.29x | 16.42x ± 24.82x | 85.4% ± 0.8% | 86.20 | 50.27 | 0.2109 ± 0.3138 |
| | Static (20%) | 3.31x ± 4.29x | 13.52x ± 19.50x | 80.9% ± 0.3% | 88.45 | 57.19 | 0.2399 ± 0.3263 |
| | Static (25%) | 3.25x ± 4.29x | 11.61x ± 16.29x | 76.6% ± 0.3% | 90.13 | 71.68 | 0.3007 ± 0.3576 |
| | Static (30%) | 3.18x ± 4.18x | 10.65x ± 15.91x | 72.3% ± 0.4% | 91.40 | 75.02 | 0.3147 ± 0.3718 |
| | **AB-NW (Ours)** | **1.16x ± 1.55x** | **8.02x ± 11.97x** | **57.7% ± 7.6%** | **99.80** | **99.26** | **0.4164 ± 0.3822** |
| **ProtBERT** | Unconstrained | 1.00x | 1.00x | 0.0% | 100.00 | 100.00 | 0.2903 ± 0.3699 |
| | Static (5%) | 2.62x ± 2.92x | 27.18x ± 27.17x | 93.2% ± 3.2% | 87.72 | 20.39 | 0.0592 ± 0.1486 |
| | Static (10%) | 2.59x ± 2.78x | 21.67x ± 26.24x | 89.7% ± 1.7% | 91.11 | 34.89 | 0.1013 ± 0.2040 |
| | Static (15%) | 2.55x ± 2.99x | 17.33x ± 20.74x | 85.4% ± 0.8% | 92.98 | 39.37 | 0.1143 ± 0.2082 |
| | Static (20%) | 2.38x ± 2.60x | 14.06x ± 16.88x | 80.9% ± 0.3% | 94.14 | 53.15 | 0.1543 ± 0.2500 |
| | Static (25%) | 2.35x ± 2.58x | 12.11x ± 14.76x | 76.6% ± 0.3% | 95.10 | 53.26 | 0.1546 ± 0.2540 |
| | Static (30%) | 2.22x ± 2.36x | 11.00x ± 13.65x | 72.3% ± 0.4% | 95.69 | 57.87 | 0.1680 ± 0.2583 |
| | **AB-NW (Ours)** | **0.89x ± 1.01x** | **8.01x ± 9.45x** | **55.3% ± 10.2%** | **99.81** | **99.52** | **0.2889 ± 0.3705** |

Table S5: Empirical alignment metrics on **Category 6: Internal Repeats** ($N$=50 pairs, average length $806 \times 806$). Values reported as Mean ± Standard Deviation.

| Model | Method | E2E Spd. | DP Spd. | Cell Sav.% | Rel Score% | Rel SP% | SP-Score |
|---|---|---|---|---|---|---|---|
| **ESM-35M** | Unconstrained | 1.00x | 1.00x | 0.0% | 100.00 | 100.00 | 0.9913 ± 0.0307 |
| | Static (5%) | 3.31x ± 2.24x | 22.24x ± 16.07x | 94.8% ± 0.9% | 100.00 | 100.00 | 0.9913 ± 0.0307 |
| | Static (10%) | 3.03x ± 2.02x | 13.30x ± 8.82x | 90.2% ± 0.1% | 100.00 | 100.00 | 0.9913 ± 0.0307 |
| | Static (15%) | 2.76x ± 2.21x | 10.06x ± 7.37x | 85.5% ± 0.2% | 100.00 | 100.00 | 0.9913 ± 0.0307 |
| | Static (20%) | 2.65x ± 1.90x | 7.89x ± 5.46x | 81.0% ± 0.1% | 100.00 | 100.00 | 0.9913 ± 0.0307 |
| | Static (25%) | 2.47x ± 1.64x | 6.71x ± 4.54x | 76.6% ± 0.1% | 100.00 | 100.00 | 0.9913 ± 0.0307 |
| | Static (30%) | 2.34x ± 1.61x | 5.89x ± 3.99x | 72.2% ± 0.1% | 100.00 | 100.00 | 0.9913 ± 0.0307 |
| | **AB-NW (Ours)** | **1.67x ± 1.07x** | **11.03x ± 8.80x** | **86.7% ± 1.6%** | **100.00** | **100.00** | **0.9913 ± 0.0307** |
| **ESM-8M** | Unconstrained | 1.00x | 1.00x | 0.0% | 100.00 | 100.00 | 0.9781 ± 0.0840 |
| | Static (5%) | 3.81x ± 1.90x | 19.93x ± 11.45x | 94.8% ± 0.9% | 100.00 | 100.00 | 0.9781 ± 0.0840 |
| | Static (10%) | 3.38x ± 1.63x | 12.07x ± 6.66x | 90.2% ± 0.1% | 100.00 | 100.00 | 0.9781 ± 0.0840 |
| | Static (15%) | 3.07x ± 1.56x | 9.04x ± 5.23x | 85.5% ± 0.2% | 100.00 | 100.00 | 0.9781 ± 0.0840 |
| | Static (20%) | 2.81x ± 1.41x | 7.10x ± 3.44x | 81.0% ± 0.1% | 100.00 | 100.00 | 0.9781 ± 0.0840 |
| | Static (25%) | 2.58x ± 1.38x | 5.95x ± 3.33x | 76.6% ± 0.1% | 100.00 | 100.00 | 0.9781 ± 0.0840 |
| | Static (30%) | 2.38x ± 1.33x | 5.19x ± 2.95x | 72.2% ± 0.1% | 100.00 | 100.00 | 0.9781 ± 0.0840 |
| | **AB-NW (Ours)** | **1.86x ± 0.93x** | **11.73x ± 6.34x** | **89.6% ± 2.2%** | **100.00** | **100.00** | **0.9781 ± 0.0840** |
| **ProtBERT** | Unconstrained | 1.00x | 1.00x | 0.0% | 100.00 | 100.00 | 0.9838 ± 0.0648 |
| | Static (5%) | 2.22x ± 1.19x | 23.67x ± 13.65x | 94.8% ± 0.9% | 100.00 | 100.00 | 0.9838 ± 0.0648 |
| | Static (10%) | 2.08x ± 1.06x | 14.40x ± 7.26x | 90.2% ± 0.1% | 100.00 | 100.00 | 0.9838 ± 0.0648 |
| | Static (15%) | 1.94x ± 0.95x | 10.72x ± 5.12x | 85.5% ± 0.2% | 100.00 | 100.00 | 0.9838 ± 0.0648 |
| | Static (20%) | 1.88x ± 0.91x | 8.42x ± 4.08x | 81.0% ± 0.1% | 100.00 | 100.00 | 0.9838 ± 0.0648 |
| | Static (25%) | 1.80x ± 0.91x | 7.14x ± 3.32x | 76.6% ± 0.1% | 100.00 | 100.00 | 0.9838 ± 0.0648 |
| | Static (30%) | 1.69x ± 0.93x | 6.26x ± 3.24x | 72.2% ± 0.1% | 100.00 | 100.00 | 0.9838 ± 0.0648 |
| | **AB-NW (Ours)** | **1.10x ± 0.60x** | **10.79x ± 5.18x** | **85.2% ± 2.1%** | **100.00** | **100.00** | **0.9838 ± 0.0648** |

Table S6: Empirical alignment metrics on **Category 7: Short Micro-Peptides** ($N$=50 pairs, average length 62 × 62). Values reported as Mean ± Standard Deviation.

| Model | Method | E2E Spd. | DP Spd. | Cell Sav.% | Rel Score% | Rel SP% | SP-Score |
|---|---|---|---|---|---|---|---|
| **ESM-35M** | Unconstrained | 1.00x | 1.00x | 0.0% | 100.00 | 100.00 | 0.9836 ± 0.0745 |
| | Static (5%) | 5.42x ± 11.16x | 55.30x ± 113.39x | 82.4% ± 3.4% | 99.80 | 96.72 | 0.9513 ± 0.2030 |
| | Static (10%) | 5.92x ± 12.49x | 55.86x ± 116.84x | 82.4% ± 3.4% | 99.80 | 96.72 | 0.9513 ± 0.2030 |
| | Static (15%) | 6.02x ± 12.59x | 58.51x ± 122.55x | 82.4% ± 3.4% | 99.80 | 96.72 | 0.9513 ± 0.2030 |
| | Static (20%) | 5.74x ± 11.88x | 49.79x ± 103.13x | 80.1% ± 1.8% | 99.80 | 96.72 | 0.9513 ± 0.2030 |
| | Static (25%) | 5.84x ± 12.04x | 51.69x ± 110.19x | 76.9% ± 0.8% | 99.80 | 96.72 | 0.9513 ± 0.2030 |
| | Static (30%) | 5.87x ± 12.59x | 45.05x ± 94.31x | 72.0% ± 0.8% | 99.82 | 97.46 | 0.9586 ± 0.1724 |
| | **AB-NW (Ours)** | **2.54x ± 5.26x** | **55.58x ± 117.22x** | **70.1% ± 5.8%** | **100.00** | **100.00** | **0.9836 ± 0.0745** |
| **ESM-8M** | Unconstrained | 1.00x | 1.00x | 0.0% | 100.00 | 100.00 | 0.9850 ± 0.0741 |
| | Static (5%) | 2.03x ± 3.61x | 16.96x ± 28.52x | 82.4% ± 3.4% | 99.70 | 96.72 | 0.9527 ± 0.2031 |
| | Static (10%) | 2.14x ± 4.25x | 17.72x ± 32.92x | 82.4% ± 3.4% | 99.70 | 96.72 | 0.9527 ± 0.2031 |
| | Static (15%) | 2.07x ± 3.96x | 18.55x ± 32.19x | 82.4% ± 3.4% | 99.70 | 96.72 | 0.9527 ± 0.2031 |
| | Static (20%) | 2.04x ± 3.80x | 16.79x ± 28.38x | 80.1% ± 1.8% | 99.70 | 96.72 | 0.9527 ± 0.2031 |
| | Static (25%) | 1.95x ± 3.90x | 14.87x ± 25.32x | 76.9% ± 0.8% | 99.70 | 96.72 | 0.9527 ± 0.2031 |
| | Static (30%) | 2.08x ± 3.96x | 14.03x ± 21.44x | 72.0% ± 0.8% | 99.71 | 96.72 | 0.9527 ± 0.2031 |
| | **AB-NW (Ours)** | **0.84x ± 1.39x** | **16.14x ± 27.59x** | **67.0% ± 6.6%** | **100.00** | **100.00** | **0.9850 ± 0.0741** |
| **ProtBERT** | Unconstrained | 1.00x | 1.00x | 0.0% | 100.00 | 100.00 | 0.9477 ± 0.2034 |
| | Static (5%) | 3.27x ± 6.81x | 39.56x ± 78.01x | 82.4% ± 3.4% | 100.00 | 100.00 | 0.9477 ± 0.2034 |
| | Static (10%) | 3.07x ± 6.07x | 37.63x ± 72.44x | 82.4% ± 3.4% | 100.00 | 100.00 | 0.9477 ± 0.2034 |
| | Static (15%) | 3.40x ± 6.79x | 40.56x ± 81.96x | 82.4% ± 3.4% | 100.00 | 100.00 | 0.9477 ± 0.2034 |
| | Static (20%) | 3.83x ± 7.60x | 38.04x ± 72.34x | 80.1% ± 1.8% | 100.00 | 100.00 | 0.9477 ± 0.2034 |
| | Static (25%) | 3.52x ± 6.76x | 33.87x ± 63.16x | 76.9% ± 0.8% | 100.00 | 100.00 | 0.9477 ± 0.2034 |
| | Static (30%) | 3.73x ± 7.38x | 31.13x ± 57.23x | 72.0% ± 0.8% | 100.00 | 100.00 | 0.9477 ± 0.2034 |
| | **AB-NW (Ours)** | **1.50x ± 3.02x** | **34.88x ± 66.68x** | **65.7% ± 6.9%** | **100.00** | **100.00** | **0.9477 ± 0.2034** |

Table S7: Empirical alignment metrics on **Category 8: Standard Benchmark Suite** ($N$=50 pairs, average length 538 × 538). Values reported as Mean ± Standard Deviation.

| Model | Method | E2E Spd. | DP Spd. | Cell Sav.% | Rel Score% | Rel SP% | SP-Score |
|---|---|---|---|---|---|---|---|
| **ESM-35M** | Unconstrained | 1.00x | 1.00x | 0.0% | 100.00 | 100.00 | 0.9735 ± 0.0832 |
| | Static (5%) | 3.64x ± 2.86x | 23.96x ± 19.17x | 95.1% ± 0.1% | 100.00 | 100.00 | 0.9735 ± 0.0832 |
| | Static (10%) | 3.32x ± 2.48x | 14.26x ± 11.48x | 90.3% ± 0.1% | 100.00 | 100.00 | 0.9735 ± 0.0832 |
| | Static (15%) | 3.01x ± 2.53x | 10.56x ± 8.73x | 85.6% ± 0.1% | 100.00 | 100.00 | 0.9735 ± 0.0832 |
| | Static (20%) | 2.85x ± 2.20x | 8.53x ± 6.88x | 81.0% ± 0.1% | 100.00 | 100.00 | 0.9735 ± 0.0832 |
| | Static (25%) | 2.67x ± 1.97x | 7.02x ± 5.27x | 76.5% ± 0.1% | 100.00 | 100.00 | 0.9735 ± 0.0832 |
| | Static (30%) | 2.53x ± 1.98x | 6.18x ± 4.89x | 72.3% ± 0.1% | 100.00 | 100.00 | 0.9735 ± 0.0832 |
| | **AB-NW (Ours)** | **1.69x ± 1.14x** | **11.26x ± 8.29x** | **86.9% ± 0.6%** | **100.00** | **100.00** | **0.9735 ± 0.0832** |
| **ESM-8M** | Unconstrained | 1.00x | 1.00x | 0.0% | 100.00 | 100.00 | 0.9819 ± 0.0632 |
| | Static (5%) | 3.36x ± 1.29x | 16.77x ± 5.28x | 95.1% ± 0.1% | 100.00 | 100.00 | 0.9819 ± 0.0632 |
| | Static (10%) | 3.01x ± 1.04x | 10.23x ± 3.26x | 90.3% ± 0.1% | 100.00 | 100.00 | 0.9819 ± 0.0632 |
| | Static (15%) | 2.67x ± 0.93x | 7.65x ± 2.49x | 85.6% ± 0.1% | 100.00 | 100.00 | 0.9819 ± 0.0632 |
| | Static (20%) | 2.48x ± 0.95x | 6.17x ± 2.04x | 81.0% ± 0.1% | 100.00 | 100.00 | 0.9819 ± 0.0632 |
| | Static (25%) | 2.29x ± 0.83x | 5.21x ± 1.74x | 76.5% ± 0.1% | 100.00 | 100.00 | 0.9819 ± 0.0632 |
| | Static (30%) | 2.15x ± 0.76x | 4.50x ± 1.47x | 72.3% ± 0.1% | 100.00 | 100.00 | 0.9819 ± 0.0632 |
| | **AB-NW (Ours)** | **1.60x ± 0.77x** | **10.13x ± 3.52x** | **89.8% ± 1.3%** | **100.00** | **100.00** | **0.9819 ± 0.0632** |
| **ProtBERT** | Unconstrained | 1.00x | 1.00x | 0.0% | 100.00 | 100.00 | 0.9945 ± 0.0173 |
| | Static (5%) | 2.51x ± 2.17x | 26.25x ± 21.71x | 95.1% ± 0.1% | 100.00 | 100.00 | 0.9945 ± 0.0173 |
| | Static (10%) | 2.37x ± 1.94x | 15.58x ± 12.76x | 90.3% ± 0.1% | 100.00 | 100.00 | 0.9945 ± 0.0173 |
| | Static (15%) | 2.20x ± 1.85x | 11.80x ± 11.39x | 85.6% ± 0.1% | 100.00 | 100.00 | 0.9945 ± 0.0173 |
| | Static (20%) | 2.13x ± 2.05x | 9.83x ± 11.52x | 81.0% ± 0.1% | 100.00 | 100.00 | 0.9945 ± 0.0173 |
| | Static (25%) | 2.04x ± 1.85x | 7.92x ± 8.09x | 76.5% ± 0.1% | 100.00 | 100.00 | 0.9945 ± 0.0173 |
| | Static (30%) | 1.94x ± 1.65x | 7.02x ± 7.03x | 72.3% ± 0.1% | 100.00 | 100.00 | 0.9945 ± 0.0173 |
| | **AB-NW (Ours)** | **1.17x ± 0.87x** | **11.73x ± 11.15x** | **85.2% ± 1.4%** | **100.00** | **100.00** | **0.9945 ± 0.0173** |

Table S8: Empirical alignment metrics on **Category 9: Unbiased Random Control** ($N$=50 pairs, average length 708 × 709). Values reported as Mean ± Standard Deviation.

| Model | Method | E2E Spd. | DP Spd. | Cell Sav.% | Rel Score% | Rel SP% | SP-Score |
|---|---|---|---|---|---|---|---|
| **ESM-35M** | Unconstrained | 1.00x | 1.00x | 0.0% | 100.00 | 100.00 | 0.9418 ± 0.2009 |
| | Static (5%) | 4.86x ± 7.14x | 33.74x ± 55.98x | 94.3% ± 2.5% | 99.68 | 101.44 | 0.9554 ± 0.1548 |
| | Static (10%) | 4.67x ± 7.37x | 23.84x ± 45.85x | 90.0% ± 1.5% | 99.90 | 102.12 | 0.9618 ± 0.1480 |
| | Static (15%) | 4.28x ± 6.79x | 19.01x ± 38.85x | 85.4% ± 0.8% | 99.92 | 100.00 | 0.9418 ± 0.2009 |
| | Static (20%) | 4.01x ± 6.57x | 14.95x ± 29.83x | 81.0% ± 0.2% | 99.92 | 100.00 | 0.9418 ± 0.2009 |
| | Static (25%) | 3.69x ± 5.97x | 12.14x ± 24.20x | 76.5% ± 0.2% | 99.92 | 100.00 | 0.9418 ± 0.2009 |
| | Static (30%) | 3.51x ± 5.87x | 11.09x ± 23.13x | 72.2% ± 0.2% | 99.95 | 100.00 | 0.9418 ± 0.2009 |
| | **AB-NW (Ours)** | **2.05x ± 2.20x** | **18.12x ± 33.26x** | **85.7% ± 3.8%** | **100.00** | **100.00** | **0.9418 ± 0.2009** |
| **ESM-8M** | Unconstrained | 1.00x | 1.00x | 0.0% | 100.00 | 100.00 | 0.9472 ± 0.1888 |
| | Static (5%) | 3.61x ± 3.27x | 20.40x ± 20.54x | 94.3% ± 2.5% | 99.80 | 99.22 | 0.9398 ± 0.1947 |
| | Static (10%) | 3.38x ± 3.21x | 13.60x ± 17.49x | 90.0% ± 1.5% | 99.99 | 100.00 | 0.9472 ± 0.1888 |
| | Static (15%) | 3.03x ± 2.82x | 10.15x ± 12.64x | 85.4% ± 0.8% | 99.99 | 100.00 | 0.9472 ± 0.1888 |
| | Static (20%) | 2.77x ± 2.45x | 8.30x ± 10.60x | 81.0% ± 0.2% | 100.00 | 100.00 | 0.9472 ± 0.1888 |
| | Static (25%) | 2.63x ± 2.24x | 6.94x ± 8.66x | 76.5% ± 0.2% | 100.00 | 100.00 | 0.9472 ± 0.1888 |
| | Static (30%) | 2.45x ± 2.45x | 6.09x ± 7.90x | 72.2% ± 0.2% | 100.00 | 100.00 | 0.9472 ± 0.1888 |
| | **AB-NW (Ours)** | **1.79x ± 1.04x** | **12.80x ± 13.65x** | **88.6% ± 5.0%** | **100.00** | **100.00** | **0.9472 ± 0.1888** |
| **ProtBERT** | Unconstrained | 1.00x | 1.00x | 0.0% | 100.00 | 100.00 | 0.9727 ± 0.1420 |
| | Static (5%) | 2.51x ± 2.73x | 26.59x ± 26.17x | 94.3% ± 2.5% | 99.93 | 99.24 | 0.9653 ± 0.1510 |
| | Static (10%) | 2.44x ± 2.73x | 17.86x ± 23.15x | 90.0% ± 1.5% | 100.00 | 100.00 | 0.9727 ± 0.1420 |
| | Static (15%) | 2.26x ± 2.46x | 13.49x ± 19.05x | 85.4% ± 0.8% | 100.00 | 100.00 | 0.9727 ± 0.1420 |
| | Static (20%) | 2.21x ± 2.25x | 11.01x ± 14.59x | 81.0% ± 0.2% | 100.00 | 100.00 | 0.9727 ± 0.1420 |
| | Static (25%) | 2.11x ± 2.23x | 9.58x ± 13.79x | 76.5% ± 0.2% | 100.00 | 100.00 | 0.9727 ± 0.1420 |
| | Static (30%) | 2.05x ± 2.52x | 8.30x ± 12.22x | 72.2% ± 0.2% | 100.00 | 100.00 | 0.9727 ± 0.1420 |
| | **AB-NW (Ours)** | **1.22x ± 1.02x** | **13.52x ± 17.74x** | **84.1% ± 4.7%** | **100.00** | **100.00** | **0.9727 ± 0.1420** |

### S2. Supplementary Benchmark Dataset Provenance and Selection Statistics

Table S9 details the candidate pool availability and exact sampled counts across the family-level split. Table S10 presents the structural distributions and source databases for all 400 held-out test sequence pairs.

Table S9: Stratified benchmark category definitions, available candidate pool sizes, and exact sampled sequence pair counts for hyperparameter tuning ($\mathcal{D}_{\text{tune}}$) and held-out testing ($\mathcal{D}_{\text{test}}$).

| | | **Tuning Set** ($\mathcal{D}_{\text{tune}}$) | | **Test Set** ($\mathcal{D}_{\text{test}}$) | |
|---|---|---|---|---|---|
| **Category** | **Selection criterion** | **Pool** | **Sampled** | **Pool** | **Sampled** |
| 1. Twilight Zone | SeqID $\leq$ 30% *and* $\text{Drift}_{\max} \geq 20$ | 1,024 | 150 | 2,339 | 50 |
| 2. Ultra-Giant Matrices | $N \times M \geq 3 \times 10^5$ cells | 46,434 | 150 | 117,550 | 25 |
| 3. Diverse Large Matrices | $N \times M \geq 3 \times 10^5$ cells, family-capped | 46,434 | 150 | 117,550 | 25 |
| 4. Asymmetric Indels | $\lvert N - M \rvert \geq 150$ (tune) / $\geq 100$ (test) | 101 | 101 | 173 | 50 |
| 5. Extreme Ratio Asymmetry | AR $\geq 2.5$ | 9 | 9 | 65 | 50 |
| 6. Internal Repeats | Keyword match or $k$-mer ($k$=4) count $\geq 8$ | 111,274 | 150 | 261,501 | 50 |
| 7. Short Micro-Peptides | $\min(N, M) \leq 80$ residues | 3,133 | 100 | 7,290 | 50 |
| 8. Standard Benchmark | Both sequences in $[200, 1000]$ residues | 54,668 | 150 | 186,676 | 50 |
| 9. Unbiased Random | All valid pairs (uniform control group) | 111,274 | 150 | 294,257 | 50 |
| **Total Disjoint Pairs** | | — | **1,110** | — | **400** |

Table S10: Aggregated structural characteristics, coarse downsampling stride ($k$), and database provenance for the held-out evaluation corpus ($\mathcal{D}_{\text{test}}$, 400 sequence pairs across 9 categories). Values reported as Mean ± SD.

| **Category / Source Databases** | **Pairs** | **Length** $N$ | **Length** $M$ | $\lvert N-M \rvert$ | **SeqID (%)** | **Drift (res)** | **Aspect Ratio** | **Stride ($k$)** |
|---|---|---|---|---|---|---|---|---|
| 1. Twilight Zone | 50 | $528 \pm 105$ | $559 \pm 123$ | $143 \pm 147$ | $20.3\% \pm 3.9\%$ | $77.2 \pm 71.3$ | $1.35 \pm 0.42$ | $10.0 \pm 1.6$ |
| BAliBASE: 40, SABmark: 6, PREFAB: 4 | | | | | | | | |
| 2. Ultra-Giant | 25 | $8481 \pm 0$ | $8481 \pm 0$ | $0 \pm 0$ | $47.9\% \pm 43.5\%$ | $0.0 \pm 0.0$ | $1.00 \pm 0.00$ | $188.0 \pm 0.0$ |
| BAliBASE: 25 | | | | | | | | |
| 3. Diverse Large | 25 | $3795 \pm 1512$ | $3795 \pm 1512$ | $0 \pm 0$ | $49.1\% \pm 29.4\%$ | $0.0 \pm 0.0$ | $1.00 \pm 0.00$ | $83.7 \pm 33.7$ |
| BAliBASE: 25 | | | | | | | | |
| 4. Asymmetric Indels | 50 | $321 \pm 181$ | $343 \pm 183$ | $243 \pm 90$ | $20.5\% \pm 16.5\%$ | $119.8 \pm 68.8$ | $2.41 \pm 0.74$ | $4.2 \pm 2.4$ |
| SABmark: 25, BAliBASE: 20, PREFAB: 5 | | | | | | | | |
| 5. Extreme Ratio | 50 | $203 \pm 170$ | $244 \pm 154$ | $226 \pm 114$ | $21.4\% \pm 19.1\%$ | $99.5 \pm 89.5$ | $3.08 \pm 0.59$ | $1.9 \pm 1.1$ |
| SABmark: 35, PREFAB: 8, BAliBASE: 7 | | | | | | | | |
| 6. Internal Repeats | 50 | $806 \pm 943$ | $806 \pm 943$ | $0 \pm 0$ | $54.7\% \pm 17.7\%$ | $0.0 \pm 0.0$ | $1.00 \pm 0.00$ | $17.5 \pm 20.9$ |
| BAliBASE: 48, OXBench: 2 | | | | | | | | |
| 7. Micro-Peptides | 50 | $62 \pm 12$ | $62 \pm 12$ | $2 \pm 4$ | $55.7\% \pm 31.2\%$ | $1.2 \pm 3.0$ | $1.03 \pm 0.08$ | $1.0 \pm 0.0$ |
| OXBench: 23, BAliBASE: 23, SABmark: 3, PREFAB: 1 | | | | | | | | |
| 8. Standard Bench | 50 | $538 \pm 226$ | $538 \pm 226$ | $0 \pm 0$ | $53.3\% \pm 17.9\%$ | $0.0 \pm 0.0$ | $1.00 \pm 0.00$ | $11.5 \pm 5.2$ |
| BAliBASE: 48, SABmark: 2 | | | | | | | | |
| 9. Unbiased Random | 50 | $708 \pm 614$ | $709 \pm 613$ | $2 \pm 8$ | $52.4\% \pm 20.2\%$ | $1.0 \pm 4.9$ | $1.01 \pm 0.05$ | $15.3 \pm 13.6$ |
| BAliBASE: 43, SABmark: 4, OXBench: 3 | | | | | | | | |

Table S11: Paired statistical tests across backbones and benchmark categories. Wilcoxon signed-rank tests ($W$) quantify differences against baseline; Cohen's $d$ reflects standardized effect sizes.

| Model | Category | Tested Metric | Mean Diff | Wilcox. $W$ | Wilcox. $p$ | Cohen's $d$ | Significant? |
|---|---|---|---|---|---|---|---|
| **ESM-35M** | 1. Twilight Zone | SP vs Static 20% | +0.1047 | 40.0 | 0.0055 | +0.3975 | **True** ($p < 0.01$) |
| | 1. Twilight Zone | DP Time vs Base | −0.0067s | 0.0 | $1.78 \times 10^{-15}$ | −2.0579 | **True** ($p < 0.001$) |
| | 2. Ultra-Giant | DP Time vs Base | −1.0090s | 0.0 | $5.96 \times 10^{-8}$ | −25.2742 | **True** ($p < 0.001$) |
| | 3. Diverse Large | DP Time vs Base | −0.2625s | 0.0 | $5.96 \times 10^{-8}$ | −1.1214 | **True** ($p < 0.001$) |
| | 4. Asym. Indels | SP vs Static 20% | +0.2643 | 0.0 | $3.68 \times 10^{-7}$ | +0.8374 | **True** ($p < 0.001$) |
| | 4. Asym. Indels | DP Time vs Base | −0.0039s | 0.0 | $1.78 \times 10^{-15}$ | −0.9021 | **True** ($p < 0.001$) |
| | 5. Extreme Ratio | SP vs Static 20% | +0.2776 | 61.0 | $3.07 \times 10^{-7}$ | +0.9072 | **True** ($p < 0.001$) |
| | 5. Extreme Ratio | DP Time vs Base | −0.0027s | 0.0 | $7.56 \times 10^{-10}$ | −0.6963 | **True** ($p < 0.001$) |
| | 6. Internal Repeats | DP Time vs Base | −0.0257s | 0.0 | $1.78 \times 10^{-15}$ | −0.4022 | **True** ($p < 0.001$) |
| | 7. Micro-Peptides | DP Time vs Base | −0.0021s | 0.0 | $7.56 \times 10^{-10}$ | −0.4717 | **True** ($p < 0.001$) |
| | 8. Standard Bench | DP Time vs Base | −0.0084s | 0.0 | $1.78 \times 10^{-15}$ | −1.2797 | **True** ($p < 0.001$) |
| | 9. Unbiased Random | DP Time vs Base | −0.0170s | 0.0 | $1.78 \times 10^{-15}$ | −0.6317 | **True** ($p < 0.001$) |
| **ESM-8M** | 1. Twilight Zone | DP Time vs Base | −0.0052s | 0.0 | $1.78 \times 10^{-15}$ | −3.0545 | **True** ($p < 0.001$) |
| | 2. Ultra-Giant | DP Time vs Base | −0.9385s | 0.0 | $5.96 \times 10^{-8}$ | −47.3491 | **True** ($p < 0.001$) |
| | 3. Diverse Large | DP Time vs Base | −0.2446s | 0.0 | $5.96 \times 10^{-8}$ | −1.1095 | **True** ($p < 0.001$) |
| | 4. Asym. Indels | SP vs Static 20% | +0.1738 | 96.5 | 0.00048 | +0.6118 | **True** ($p < 0.001$) |
| | 5. Extreme Ratio | SP vs Static 20% | +0.1765 | 167.5 | $5.91 \times 10^{-5}$ | +0.6385 | **True** ($p < 0.001$) |
| | 6. Internal Repeats | DP Time vs Base | −0.0234s | 0.0 | $1.78 \times 10^{-15}$ | −0.3941 | **True** ($p < 0.001$) |
| | 7. Micro-Peptides | DP Time vs Base | −0.0007s | 0.0 | $7.55 \times 10^{-10}$ | −0.5080 | **True** ($p < 0.001$) |
| | 8. Standard Bench | DP Time vs Base | −0.0073s | 0.0 | $1.78 \times 10^{-15}$ | −1.3486 | **True** ($p < 0.001$) |
| | 9. Unbiased Random | DP Time vs Base | −0.0167s | 0.0 | $1.78 \times 10^{-15}$ | −0.5731 | **True** ($p < 0.001$) |
| **ProtBERT** | 1. Twilight Zone | SP vs Static 20% | +0.0329 | 0.0 | 0.0254 | +0.2413 | **True** ($p < 0.05$) |
| | 1. Twilight Zone | DP Time vs Base | −0.0093s | 0.0 | $1.78 \times 10^{-15}$ | −1.8567 | **True** ($p < 0.001$) |
| | 2. Ultra-Giant | DP Time vs Base | −1.3157s | 0.0 | $5.96 \times 10^{-8}$ | −31.1170 | **True** ($p < 0.001$) |
| | 3. Diverse Large | DP Time vs Base | −0.3556s | 0.0 | $5.96 \times 10^{-8}$ | −1.0862 | **True** ($p < 0.001$) |
| | 4. Asym. Indels | SP vs Static 20% | +0.1633 | 73.0 | 0.00045 | +0.5182 | **True** ($p < 0.001$) |
| | 4. Asym. Indels | DP Time vs Base | −0.0046s | 0.0 | $1.78 \times 10^{-15}$ | −0.8605 | **True** ($p < 0.001$) |
| | 5. Extreme Ratio | SP vs Static 20% | +0.1346 | 37.5 | 0.00015 | +0.5055 | **True** ($p < 0.001$) |
| | 5. Extreme Ratio | DP Time vs Base | −0.0027s | 0.0 | $1.78 \times 10^{-15}$ | −0.6967 | **True** ($p < 0.001$) |
| | 6. Internal Repeats | DP Time vs Base | −0.0315s | 0.0 | $1.78 \times 10^{-15}$ | −0.3829 | **True** ($p < 0.001$) |
| | 7. Micro-Peptides | DP Time vs Base | −0.0017s | 0.0 | $7.55 \times 10^{-10}$ | −0.4609 | **True** ($p < 0.001$) |
| | 8. Standard Bench | DP Time vs Base | −0.0090s | 0.0 | $1.78 \times 10^{-15}$ | −1.5340 | **True** ($p < 0.001$) |
| | 9. Unbiased Random | DP Time vs Base | −0.0200s | 0.0 | $1.78 \times 10^{-15}$ | −0.5732 | **True** ($p < 0.001$) |

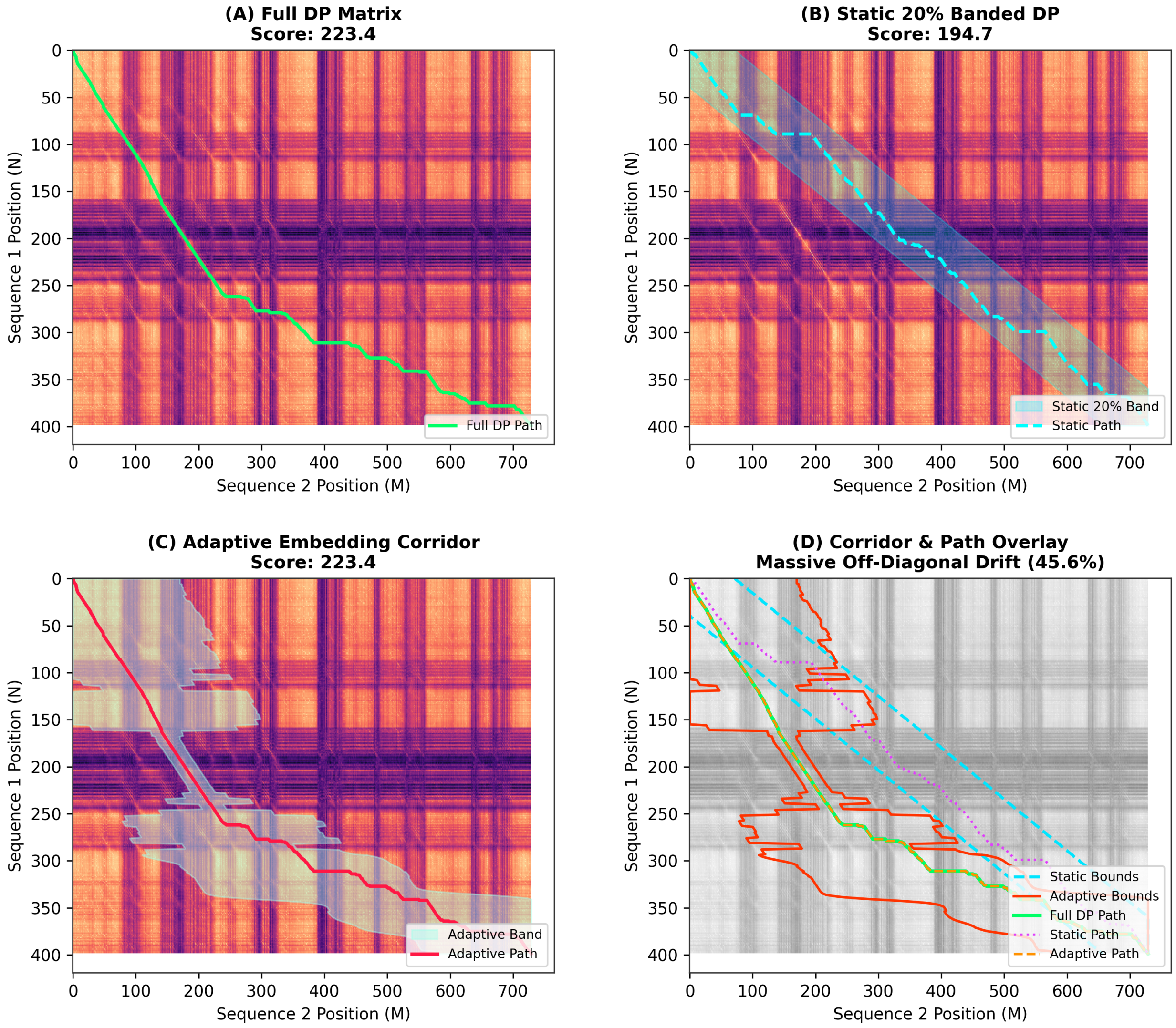


Figure S1: Global alignment decoding on pair `1oyc_2tmdA` exhibiting 45.6% off-diagonal drift from Category 1 ($400 \times 730$ footprint). **(A)** Unconstrained DP matrix evaluating all $\mathcal{O}(NM)$ cells (optimal score: 223.4). **(B)** Static 20% banded DP corridor clips the drifting path, dropping score to 194.7. **(C)** Adaptive corridor tracks the off-diagonal trajectory, retaining the exact 223.4 score. **(D)** Bound and path overlay: static bounds (cyan dotted), adaptive bounds (red solid), full DP path (green), static path (magenta dotted), adaptive path (yellow dashed).

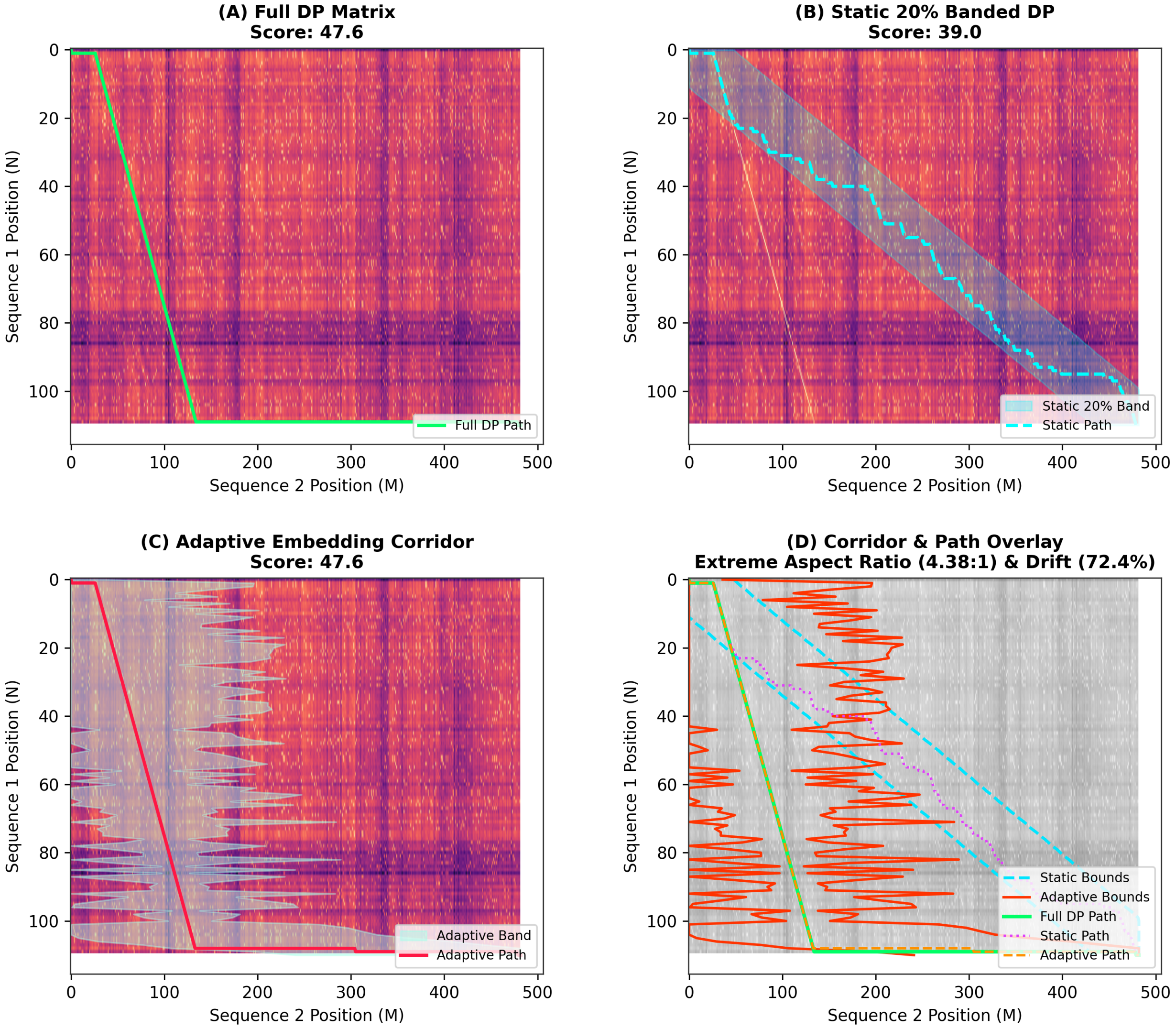


Figure S2: Alignment on asymmetric pair `1krs_1lylA` (4.38:1 aspect ratio, 72.4% drift, 110 × 480 matrix) from Category 5. **(A)** Unconstrained DP matrix (score: 47.6). **(B)** Static 20% band clips the trajectory along the diagonal, lowering score to 39.0. **(C)** Adaptive corridor adjusts row span $w_i$, recovering baseline score 47.6. **(D)** Bound overlay showing adaptive corridor (red) containing the full path (green).
